# Cardiac electrical abnormalities in a mouse model of left ventricular non-compaction cardiomyopathy

**DOI:** 10.1101/2024.11.20.624427

**Authors:** Vítor S. Fernandes, Ricardo Caballero, Marcos Siguero-Álvarez, Tania Papoutsi, Juan Ramón Gimeno, Eva Delpón, José Luís de la Pompa

**Affiliations:** Intercellular Signaling in Cardiovascular Development and Disease Laboratory, Centro Nacional de Investigaciones Cardiovasculares Carlos III (CNIC), Calle Melchor Fernández Almagro 3, 28029 Madrid, SPAIN; Ciber de Enfermedades Cardiovasculares, Instituto de Salud Carlos III, Calle Melchor Fernández Almagro 3, 28029 Madrid, SPAIN; Departamento de Farmacología, Facultad de Medicina, Universidad Complutense de Madrid, Plaza Ramón y Cajal S/N, 28040 Madrid, SPAIN; Unidad CSUR de Cardiopatías Familiares, Servicio de Cardiología, Hospital Universitario Virgen de la Arrixaca, Universidad de Murcia, El Palmar, 30120 Murcia, SPAIN

## Abstract

Mutations in *MINDBOMB 1* (*MIB1*), encoding an E3 ubiquitin ligase of the NOTCH signaling pathway, cause left ventricular noncompaction cardiomyopathy (LVNC) in mice and humans, increasing the risk of arrhythmia and left ventricular dysfunction. This study aimed to investigate the effect of *MIB1* mutations on cardiac electrical activity. We examined male *Mib1^flox^;Tnnt2^Cre^* mice, a disease model of LVNC, and wildtype littermates on the C57BL/6J genetic background. Our results demonstrate that the gap-junction protein connexin43 was delocalized from the intercalated disks to the lateral long axis of *Mib1^flox^;Tnnt2^Cre^* cardiomyocytes. Cardiomyocyte electrophysiology revealed an increase in the Na (I_Na_) peak density at potentials between –50 and –30 mV in *Mib1^flox^;Tnnt2^Cre^*mice, with no changes in I_Na_ activation or inactivation kinetics. *Mib1^flox^;Tnnt2^Cre^* cardiomyocytes also showed decreases in outward K^+^ peak currents and currents at the end of depolarizing pulses at potentials ≥−10 mV and ≥−20 mV, respectively, and this was accompanied by a lower charge density at ≥−20 mV. Action potential duration was increased in *Mib1^flox^;Tnnt2^Cre^* cardiomyocytes. The cardiac stress, induced by swimming endurance training or β-adrenergic stimulation with isoproterenol, increases QTc duration in *Mib1^flox^;Tnnt2^Cre^* mice, accompanied by a decrease in T-wave amplitude and area. Swimming endurance training decreased heart rate in wildtype and *Mib1^flox^;Tnnt2^Cre^* mice but was unaffected by long-term isoproterenol treatment. These mouse findings are in agreement with an increased QTc duration found in LVNC patients carrying *MIB1* mutations. These results provide insight into the outcomes of LVNC and relate its pathogenicity to impaired ventricular repolarization.

## Introduction

Left ventricular non-compaction (LVNC) is a heterogeneous cardiomyopathy with a poorly understood etiology and is believed to result from *in utero* alteration of ventricular chamber development [1], although isolated cases of acquired and potentially reversible LVNC have also been reported [2]. LVNC is characterized by prominent trabeculations, deep endocardial recesses in the ventricular wall, and a thin compact myocardium layer [3]. The clinical consequences of these structural abnormalities vary, ranging from mild effects with normal heart function to congestive heart failure, cardiac conduction abnormalities, ventricular tachyarrhythmia, and sudden cardiac death [3–6]. Ventricular arrhythmia and myocardial dysfunction in LVNC patients are predictors of premature death [6].

LVNC is genetically heterogeneous, with a predominantly autosomal-dominant inheritance pattern [7], and has been linked to sarcomere gene mutations, particularly in *MYH 7* [7, 8]. Other mutations implicated in LVNC affect genes encoding scaffold and nuclear proteins [8, 9]. The proposed developmental origin of LVNC suggests the involvement of genetic alterations affecting the signals and transcription factors that regulate cardiovascular development [10–13]. A key mediator of cell fate specification and tissue patterning in metazoans is the highly conserved signaling pathway NOTCH [14, 15], whose disruption in humans leads to developmental abnormalities affecting the heart and vessels [16–19]. Functional studies in mice have shown that NOTCH is crucial for the endocardium-to-myocardium signaling that governs the development of the cardiac valves and ventricular chambers and have shed light on the disease mechanisms associated with NOTCH dysfunction [reviewed in [20, 21]]. We previously showed that LVNC in mice and humans can be caused by mutations in the gene encoding the ubiquitin ligase MINDBOMB1 (MIB1) [22], which is required for NOTCH ligand endocytosis and signaling activation [23]. Myocardium-specific inactivation of *Mib1* in mice disrupts ventricular maturation and patterning, so that persistent trabeculae and thinner ventricular walls are found in the adult heart, severely impairing its function [22, 24, 25]. LVNC families carrying disease-co-segregating germline *MIB1* mutations have large persistent trabeculae and severely reduced ventricular function[22]. LVNC is usually associated with abnormal electrocardiographic (ECG) patterns, such as intraventricular conduction delays, voltage signs of left ventricular hypertrophy, left axis deviation, repolarization abnormalities, and QTc prolongation[26, 27]. The aim of this study was to investigate the mechanisms underlying ECG alterations associated with LVNC, utilizing a previously described mouse model of this condition.

## Material and Methods

### Animal studies

Animal studies were approved by the CNIC Animal Experimentation Ethics Committee, the Complutense University of Madrid and the Community of Madrid (Ref. PROEX 155.7/20). All animal procedures conformed to EU Directive 2010/63EU and Recommendation 2007/526/EC regarding the protection of animals used for experimental and other scientific purposes, enacted in Spanish law under Real Decreto 1201/2005. We used 20-week-old male *Mib1^flox/flox^;Tnnt2^Cre/+^* (hereafter *Mib1^flox^;Tnnt2^Cre^*) mutants and their respective wildtype littermates *Mib1^flox/flox^;Tnnt2 ^+/+^* (WT), congenic on the C57BL/6J background. Mice were housed in wire-bottomed cages in a temperature-controlled room (22±0.8°C) with a 12 h light–dark cycle and a relative humidity of 55±10%. Mice had free access to food and water.

### Mouse electrocardiography

ECGs were recorded in anesthetized mice as previously described [28]. Briefly, mice were placed in a supine position on a temperature-controlled surface and were anesthetized with 1.5-2% isoflurane. Four needle electrodes were subcutaneously inserted into the limbs. ECG recordings were acquired with a MP36R Biopac System and analyzed offline with LabChart7 (AD Instruments, Australia). A high-pass filter setting of 0.5 Hz was applied to remove the baseline wander using a bidirectional filtering strategy. The PR interval was measured from the beginning of the positive deflection of the P wave to the peak of the R wave, the QRS interval was measured from the start of the Q wave to the point where the S wave crosses the isoelectric line, and the QT interval was measured from the start of the Q wave to the point where the T wave reaches the 90% of the decline[28]. All these intervals were calculated as the mean of ∼ 450 ECG measurements, and QTc duration was calculated by considering the correction for RR intervals according to Bazett’s formula for murine ECG[29]:

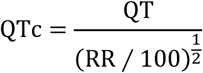

The T wave amplitude was measured as the T peak in mV and the area of the T wave was measured as the area under the curve averaging 5 T waves areas from each animal.

### Human electrocardiography

ECGs from patients with LVNC were obtained in accordance with human subject guidelines and following the approved protocol by the Hospital Universitario Virgen de la Arrixaca, Murcia, Spain. All patients provided written informed consent. Human ECG measurements were recorded using standard methods in patients carrying the *MIB1* mutations and in healthy relatives to assess differences in ventricular repolarization. Lead II and precordial V5 traces, commonly used to diagnose repolarization abnormalities^3^, were selected to assess temporal and morphological differences between patients and healthy relatives. QTc duration was calculated by considering the correction for RR intervals according to Bazett’s. The following short-term QT variability markers were calculated according to the formulas listed below: QT variance (QTvar), standard deviation of the QT intervals (SDqt), short-term variability of the QT intervals (STVqt), QT variance normalized to mean QT interval (QTVN), QT variability index (QTVI_HR_), and the root mean square of the successive QT interval differences (RMSSDqt).

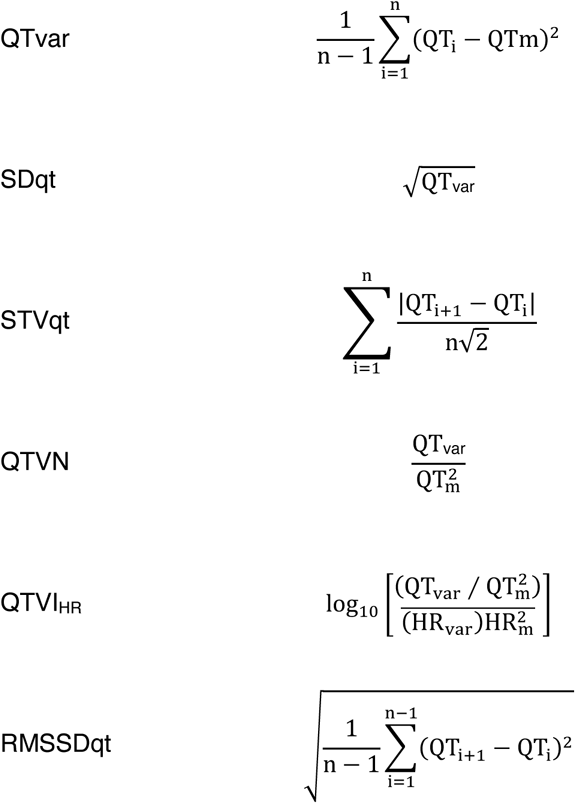

### Mouse endurance training

The endurance training protocol was as described previously [30]. Briefly, mice were first acclimatized to water by being placed in glass container (60×30×45 cm) with shallow water at a constant warm temperature (∼30°C) to minimize environmental stress. The endurance training took place during the dark phase of the light-dark cycle, corresponding to peak mouse activity. After the adaption period, the mice gained swimming experience over 2 weeks, with the swimming duration increasing by 5 min per day until reaching 45 min at the end of the second week. Over the next 3 weeks (weeks 3 to 6), the swimming duration increased by 15 min each week and was then maintained at 90 min per day during weeks 6 to 8 (**S1 Table**).

### Isoproterenol treatment

Isoproterenol is a synthetic non-selective β1– and β2-adrenergic receptor agonist that is widely used to measure the ability of the heart to respond to stress stimuli[31]. WT and *Mib1^flox^;Tnnt2^Cre^* mice received twice-weekly i.p. bolus injections of isoproterenol (3 mg/kg) over a period of 4 weeks. ECGs were recorded at baseline (before the first isoproterenol injection), and at the end of the isoproterenol administration protocol. Data were recorded and analyzed as described in **§Mouse electrocardiography**.

### Blood pressure measurements

Blood pressure was measured with an automated tail-cuff device (Visitech System BP2000, NC). Mice received training for one week to adapt to the system. Weekly measurements were then taken at the same time of day. The first 10 measurements on each measurement day were rejected, and results are presented as the mean of the last 10 measurements.

### Single cell electrophysiology

*Recording techniques*: Single ventricular myocytes were isolated from cardiac tissue by enzymatic dissociation with collagenase type II (Worthington Biochemical Corporation Lakewood, NJ, USA) and protease (type XIV, Sigma Chemical Co. London, UK)[32, 33]. Mice were heparinized (5000 U/kg i.p.) and anesthetized with ketamine (150 mg/kg i.p.) and xylazine (10 mg/kg i.p.). Currents were recorded at room temperature (21-23°C) by whole-cell patch clamping using an Axopatch-200B patch clamp amplifier (Molecular Devices, USA)[32, 33]. Recording pipettes were pulled from 1.0 mm o.d. borosilicate capillary tubes (GD1, Narishige Co., Ltd, Japan) using a programmable patch micropipette puller (Model P-2000 Brown-Flaming, Sutter Instruments Co., USA) and were heat-polished with a microforge (Model MF-83, Narishige). For measurements of micropipette resistance, the micropipette was filled with the internal solution and immersed in the external solution. Micropipette resistance was kept below 1.5 MΩ for the sodium currents (I_Na_), below 3.5 MΩ for other currents, or above 7 MΩ for measurement of action potentials (AP). In all experiments, series resistance was compensated manually by using the series resistance compensation unit of the Axopatch amplifier, and ≥80% compensation was achieved. Uncompensated access resistance was 2.2±1.8 MΩ (n=133). Thus, considering the myocyte capacitance, we expected no significant voltage errors (<5 mV) due to series resistance with the micropipettes used. To minimize the contribution of time-dependent shifts in channel availability during I_Na_ recordings, all data were collected 20 min after establishing the whole-cell configuration. Under these conditions, current amplitudes and voltage dependence of activation and inactivation were stable during recordings [32, 33]. The current recordings were sampled at 4 kHz (except for I_Na_, which was sampled at 50 kHz), filtered at half the sampling frequency, and stored on a computer hard disk for subsequent analysis.

*Solutions*: For recordings of I_Na_, the external solution contained (mM): NaCl 4, MgCl_2_ 1.0, CaCl_2_ 1.0, CdCl_2_ 1.0, CsCl1 33.5, HEPES 20, and glucose 11 (pH=7.35 with CsOH). Recording pipettes were filled with an internal solution containing (mM): NaF 10, CsF 110, CsCl 20, HEPES 10, and EGTA 10 (pH 7.35 with CsOH). The L-type Ca^+^ currents (I_CaL_) were recorded in myocytes superfused with an external solution containing (mM): tetraethylammonium 137, CaCl_2_ 1, MgCl_2_ 0.5, HEPES 10, and glucose 10 (pH=7.4 with CsOH). The internal solution contained (mM): CsCl 125, TEA-Cl 20, MgATP 5, phosphocreatine 3.6, HEPES 10, and EGTA 10 (pH=7.2 with CsOH). For recordings of inward rectifier currents (I_K1_), the external solution contained (mM): NaCl 140, 4 mM KCl, CaCl_2_ 1, MgCl_2_ 1, HEPES 10, 4-aminopyridine 2, glucose 10, and nifedipine 1 µM, atropine 0.1 µM, and glibenclamide 10 µM (pH=7.4 with NaOH). Recording pipettes were filled with an internal solution containing (mM): K-aspartate 80, KCl 42, KH_2_PO_4_ 10, MgATP 5, phosphocreatine 3, HEPES 5, and EGTA 5 (pH 7.2 with KOH). For outward K^+^ currents, the external and internal solutions were identical to those used for I_K1_ recordings, but without 4-aminopyridine in the external solution. For AP recordings, the external solution contained (mM): NaCl 136, KCl 4, CaCl_2_ 1.8, MgCl_2_ 1, HEPES 10, and glucose 10 (pH 7.4 with NaOH). The internal solution contained (mM): K-aspartate 80, KCl 42, KH_2_PO_4_ 10, MgATP 5, phosphocreatine 3, HEPES 5, and EGTA 5 (pH=7.2 with KOH). The currents were not corrected for the liquid junction potentials.

*Pulse protocols*: To construct the current-voltage relationships for I_Na_, we applied 50-ms pulses in 5 mV increments from −120 mV to potentials between −100 and +30 mV. I_Na_ amplitude was measured as the difference between the peak transient current and the current at the end of the pulses and was normalized to cell capacitance to obtain the density. To construct the inactivation curves, I_Na_ was recorded by applying 500-ms pulses from −120 mV to potentials between −140 and –20 mV in 10 mV increments followed by a test pulse to −20 mV. To analyze the recovery from inactivation of I_Na_, we applied two 50-ms pulses (P1 and P2) from –120 to −40 mV at increasing coupling intervals (0.05-500 ms). Reactivation kinetics were measured by fitting a mono exponential function to the data. The late component of I_Na_ (I_NaL_) was recorded by applying 500-ms pulses from –120 to –40 mV, and was measured as the percentage of the peak I_Na_ recorded. To construct current-voltage relationships for I_CaL_, we applied 500-ms pulses in 5 mV increments from –80 mV to potentials between –40 and +50 mV. The I_Na_ was inactivated by applying a 50-ms prepulse to –30 mV. The L-type calcium current (I_CaL_) was measured as the difference between the peak current and the current at the end of the pulses and was normalized to the cell capacitance to calculate the I_CaL_ density. Inactivation curves for I_CaL_ were obtained by applying a 500-ms conditioning pulse from –70 mV to potentials between –90 and +50 mV, followed by a test pulse to 0 mV. Conductance-voltage curves for I_Na_ and I_CaL_ were constructed by plotting the normalized conductance as a function of the membrane potential. The conductance was estimated for each experiment using the equation listed below: *G* is the conductance at the test potential V_m_, *I* represent the peak maximum current at V_m_, and E_rev_ is the reversal potential.

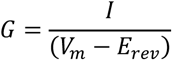

To determine the E_rev_, I_Na_ and I_CaL_ density-voltage relationships obtained in each experiment were fitted to a function of the form below: where *I* is the peak current elicited at the test potential V_m_, G_max_ is the maximum conductance, and *k* is the slope factor.

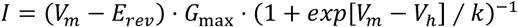

The protocol to record I_K1_ consisted of 250-ms steps imposed in 10 mV increments from –80 mV to potentials ranging –120 and –30 mV. I_K1_ was always measured at the end of the 250-ms pulses. Current–voltage relationships for outward K^+^ currents were constructed by applying 500-ms pulses in 10 mV increments from –80 mV to potentials ranging –80 and +50 mV. Peak outward K^+^ currents were measured as the difference of the peak and the current at the end of the pulse, while sustained outward K^+^ currents were measured at the end of the pulses. The charge was estimated from the integral of the current traces comprising the area between the peak and the current at the end of the 500-ms pulse. APs elicited by depolarizing-current pulses of 2 ms in duration at 1.5-2 times the current threshold at a frequency if 1Hz were recorded in the current-clamp configuration of the patch-clamp technique.

### Immunohistochemistry

Hearts from WT and *Mib1^flox^;Tnnt2^Cre^* mice were fixed overnight at 4°C in 4% paraformaldehyde in phosphate saline buffer (PBS), pH 7.4. Hearts were then dehydrated through a graded series of ethanol concentrations. The tissue was then embedded in paraffin, and longitudinal 5 μm sections were cut on a microtome. The sections were dewaxed and cleaned with PBS. Antigens were retrieved by heating the sections in sodium citrate buffer (10 mM). Endogenous biotin was blocked using an avidin/biotin kit (Vector Laboratories), and non-specific IgG binding sites were blocked with histoblock solution (5% goat serum, 3% BSA 20 mM MgCl_2_, and 0.3% Tween 20). Sections were then incubated overnight at 4°C with anti-CX43 (1:200, Sigma-Aldrich) and anti-N-cadherin (1:100, Invitrogen, Life Technologies). After several washes in PBS, samples were incubated for 2h with biotinylated anti-rabbit IgG (1:200, Vector Laboratories), washed again, and incubated for 2h at room temperature with TRITC-coupled extravidin and Alexa Fluor 488 (1:200, Thermo Fisher Scientific). Cell membranes were stained with wheat germ agglutinin (WGA Rhodamine, 1:100, Vector Laboratories). Cell nuclei were stained with DAPI (1:1000, Sigma-Aldrich). For immunofluorescence of ion channels, single ventricular myocytes were isolated from hearts by enzymatic dissociation (liberase blendzyme, Roche) as previously described[34]. Briefly, hearts were excised and retrogradely perfused at 37°C for 6–8 min with a modified tyrode solution (113 mM NaCl, 4.7 mM KCl, 0.6 mM KH_2_PO_4_, 0.6 mM Na_2_HPO_4_, 1.2 mM MgSO_4_-7H_2_O, 0.032 mM Phenol red, 12 mM NaHCO_3_, 10 mM KHCO_3_, 10 mM 4-(2-hydroxyethyl)-1-piperazineethanesulfonicacid, and 30 mM taurine, pH 7.4). Next, hearts were perfused with the same buffer containing 0.1 mg/ml liberase (Roche) until the complete digestion was achieved. The ventricles were then mechanically dissociated and single cells were transferred to a test tube containing the same solution but with the enzyme replaced with bovine calf serum (BCS, 5%) and progressive concentration of CaCl_2_ to reach a final concentration of 1 mM. Cells were fixed in 4% paraformaldehyde in PBS for 30 min at room temperature and then washed three times for 5 min with PBS containing 2% BSA. The cells were then permeabilized with 1% Triton X-100 at 25 °C for 10 min. Non-specific IgG binding sites were blocked with histoblock solution. Cells were rinsed repeatedly and incubated overnight at 4°C with the primary antibodies anti-K_V_4.2 (1:200, Alomone labs, Israel), anti-K_V_4.3 (1:200, Alomone labs, Israel), anti-Na_V_1.5 (1:200, Alomone labs, Israel), and anti-α-actinin (1:200, Sigma). All image analysis were performed using Fiji (ImageJ, NIH).

### Picrosirius red histology

Heart sections were obtained as described in immunohistochemistry section. Briefly, the nuclei were stained with Weigert’s haematoxilin for 8 min, and then the slides were washed for 10 min in running tap water. Next, slides were stained in picrosirius red solution for 1 h and then washed in two changes of acidified water. The slides were then dehydrated in 3 changes of 100% ethanol and cleared in xylene. The slides were mounted in a resinous medium. All images were analyzed using Fiji (ImageJ, NIH) to determine collagen area and data were collected from 3 section per animal.

#### Western Blot

Hearts from WT and *Mib1^flox^;Tnnt2^Cre^* mice were obtained immediately harvested after mice were sacrificed and frozen in isopentane, and stored at –80°C to carry out biochemical studies. Protein samples (20-40 µg) from each frozen tissue were denatured and resolved using SDS-PAGE (sodium dodecyl-sulfate polyacrylamide gel electrophoresis). The separated proteins were transferred to a PVDF membrane (GE Healthcare, UK). To block non-specific binding, the membranes were incubated for one hour at room temperature in TBST containing 5% nonfat dried milk. Following this, the membranes were incubated with primary antibodies: anti-KV4.2 (1:200, Alomone Labs, Israel), anti-KV4.3 (1:400, Alomone Labs, Israel), and anti-NaV1.5 (1:100, Alomone Labs, Israel). The membranes were then incubated with HRP-conjugated secondary antibodies (1:5000; GE Healthcare, UK) for 2 hours at room temperature. The peroxidase activity was detected using an Enhanced Chemiluminescence (ECL) Kit (GE Healthcare, Buckinghamshire, UK), and the signals were visualized with a Gel-Doc system using Quantity One 4.5.1 software (Bio-Rad, Hercules, USA). For loading normalization, stain-free technology was applied (Bio-Rad, Spain), and images of both the target proteins and the total protein were analyzed using ImageLab software (version 6.0.0, Bio-Rad, Hercules, USA) to determine the intensity per mm² of each band and lane. The relative protein levels were normalized to the intensity of GAPDH loading blots.

### Statistical analysis

Results are presented as the mean±SD *n* animals or mean±SEM of *n* experiments as indicated. Differences were analyzed by parametric unpaired two-tailed *t-test* or the non-parametric analog Mann–Whitney or Wilcoxon’s test for comparisons between two groups and by ANOVA followed by the Tukey or Bonferroni post-hoc test for multiple comparisons. To consider repeated sample assessments, data were analyzed with multilevel mixed-effects models. Comparisons between categorical variables were by the Fisher exact test. Differences were considered statistically significant at *p<0.05*.

## Results

### Mutations in the human *MIB1* homolog are associated with long QT prolongation

*MIB1* mutations were previously reported in two LVNC families [22]. ECG comparisons between affected individuals carrying the MIB1^V943F^ or MIB1^R530X^ mutations and their healthy relatives (**S1 Fig.**) showed no significant differences in heart rate, PR interval, QRS duration, or RR interval (**Fig. 1A,B**; **S1 Table**). However, those with *MIB1* mutations exhibited longer corrected QT intervals (QTc) compared to healthy relatives ((**Fig. 1C**; **S1 Table**, 493.5±9.1 ms vs 430.8±27.5 ms). Notably, some mutation carriers displayed left axis deviation (**S1 Table**). Analysis of QT variability markers revealed elevated values in *MIB1* mutation carriers, indicating an increased risk of ventricular arrhythmias (**Fig. 1D-I** and **S2 Table).** These results suggest that individuals carrying *MIB1* mutations have increased markers of QT variability and thus an elevated risk of ventricular arrhythmias.

**Figure 1.**
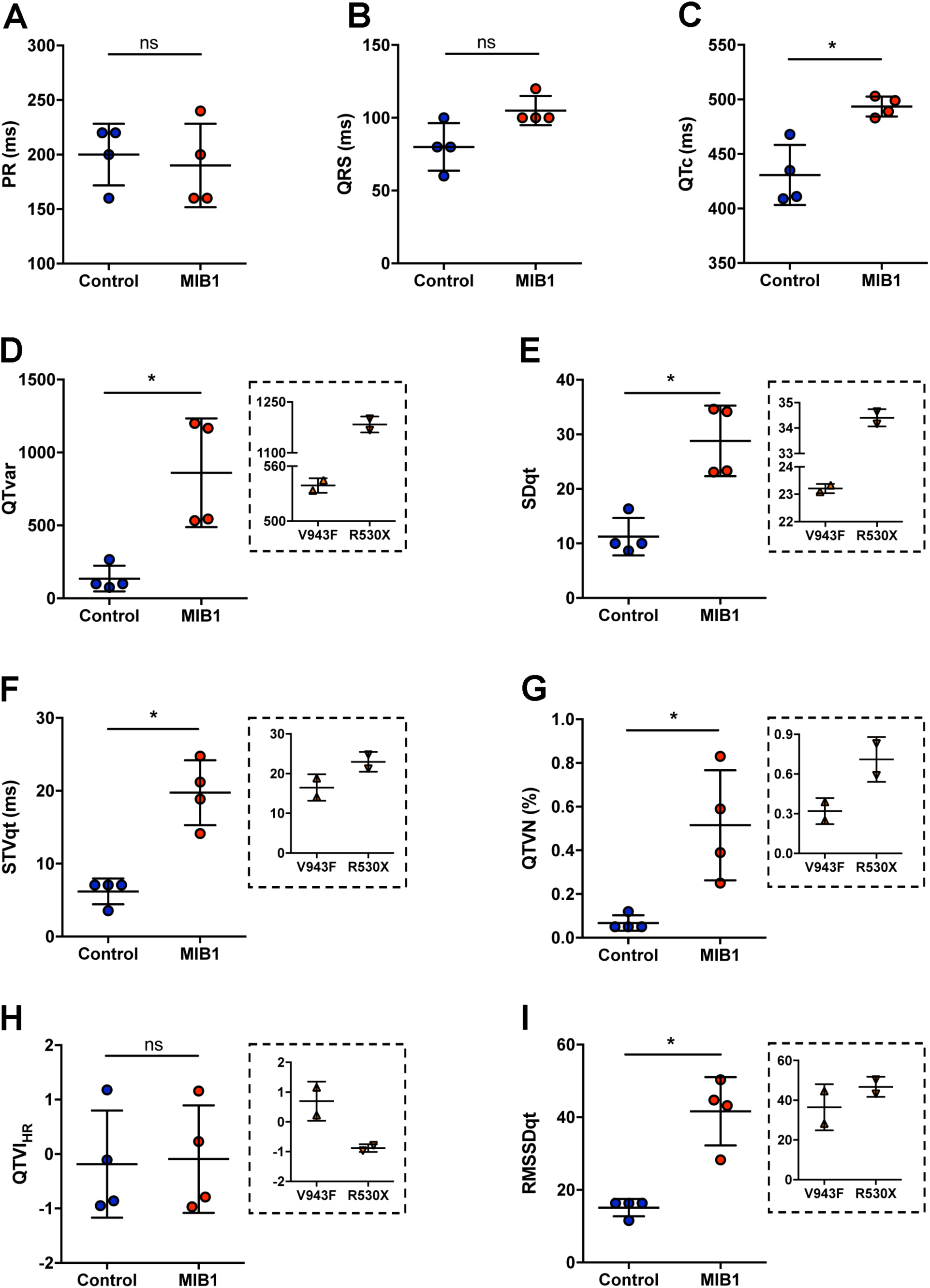
ECG abnormalities in LVNC patients carrying *MIB1* mutations. (A) PR interval measurements in healthy family members (control) and LVNC patients. (B) QRS measurements in healthy family members (control) and LVNC patients. (C) QTc measurements using the Bazzet formula for healthy family members (control) and LVNC patients. (D-I) Short-term QT variability markers predicting arrhythmias and sudden cardiac death in healthy family members (control) and LVNC patients: (D) QT variance, (E) standard deviation of QT intervals, (F) short-term beat-to-beat temporal QT variability, (G) QT variance normalized to mean QT interval, (H) QT variability index (log ratio of normalized QT variance to normalized heart rate variance), and (I) root mean square of successive QT interval differences. Insets show QT variability markers for LVNC patients with V943F or R530X *MIB1* mutations. Statistical significance was determined by the Mann–Whitney test. Results are mean±SD of 4 patients. ns, non-significant. *\*P* < 0.05 vs control.

### ECG abnormalities and prolonged action potential duration in *Mib1^flox^;Tnnt2^Cre^* mice

We investigated the electrocardiographic phenotype of *Mib1^flox^;Tnnt2^Cre^* mice, a model of LVNC due to impaired NOTCH signaling [22], to compare it with patients carrying *MIB1* mutations. While these mice showed reduced ejection fraction at 6 months[22], there were no differences in systolic or diastolic pressure between wild type (WT) and mutant mice, both at baseline and after swimming endurance training (**S2A,B Fig.**). Heart weight/tibial ratio was also similar between groups (**S2C Fig.**). Baseline measurements of heart rate (**Fig. 2A,B,C**), PQ interval (**Fig. 2D,G**), QRS duration (**Fig. 2E,H**), and QT_c90_ interval (**Fig. 2F,I**), revealed no differences between WT and *Mib1^flox^;Tnnt2^Cre^* mice. An 8-week swimming regimen (**S3 Table**), during which both WT and *Mib1^flox^;Tnnt2^Cre^* mice swam similar distances (**S2D Fig. 2** and **S1,2 Videos**) resulted in a slight but significant decrease in heart rate in WT mice, with no difference observed between WT and *Mib1^flox^;Tnnt2^Cre^* groups (**Fig. 2B**). One WT mouse and four *Mib1^flox^;Tnnt2^Cre^* mice died during the training (**S2E Fig.**). Additionally, swimming endurance training did not significantly affect the PQ interval (**Fig. 2D**). However, *Mib1^flox^;Tnnt2^Cre^* mutants exhibited increased QRS (**Fig. 2E**) and QT_c90_ intervals (**Fig. 2F**) compared to WT, indicating exercise-induced conduction abnormalities.

**Figure 2.**
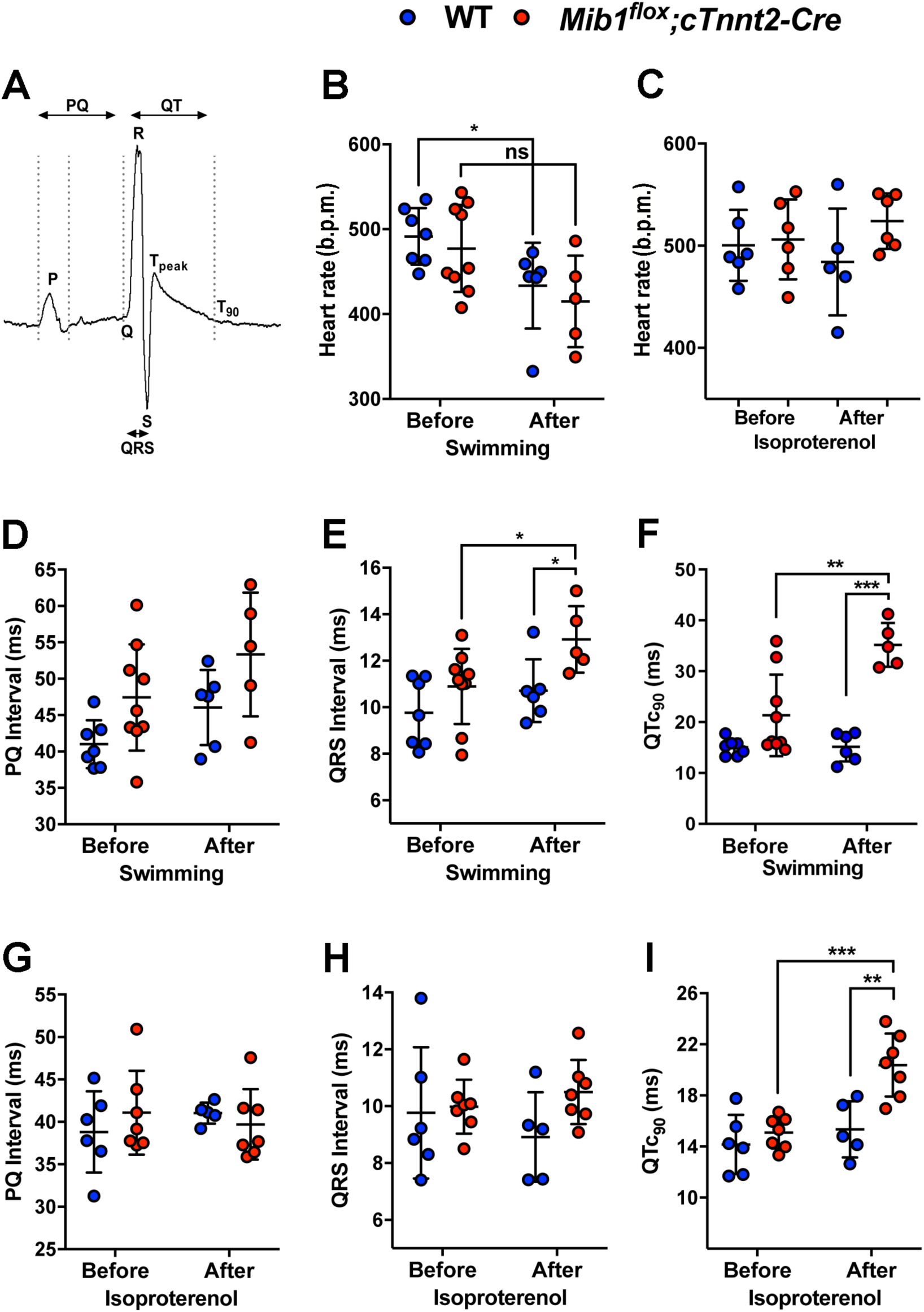
ECG alterations in *Mib1^flox^;Tnnt2^Cre^* mice are associated with cardiac stress from isoproterenol or swimming endurance training. (A) ECG recording showing measured intervals. (B, C) Mean heart rate (bpm) of isoflurane-anesthetized WT and *Mib1^flox^;Tnnt2^Cre^* mice before and after swimming endurance training (B) or at the end of the isoproterenol protocol (C). (D) PQ, (E) QRS, and (F) QTc90 duration measured before and after swimming endurance training. (G) PQ, (H) QRS, and (I) QTc90 duration measured before and at the end of the isoproterenol protocol. Statistical significance was determined by ANOVA with Tukey post-hoc test. *\*P* < 0.05 vs WT. Results are mean±SD of 6-9 animals.

Chronic isoproterenol treatment showed no differences in heart rate (**Fig. 2C**), PQ interval (**Fig. 2G**) or QRS duration (**Fig. 2H**) between genotypes, but QT_c90_ was significantly prolonged in *Mib1^flox^;Tnnt2^Cre^* mutants (**Fig. 2I**). T-wave amplitude and area under the curve were comparable at baseline (**Fig. 3A,A’**), but decreased significantly in *Mib1^flox^;Tnnt2^Cre^* mice after swimming (**Fig. 3B,B’,C,D**), suggesting exercise-induced ventricular repolarization abnormalities. Next, we recorded action potentials (APs) from cardiomyocytes dissociated from *Mib1^flox^;Tnnt2^Cre^* mice, paced at 1 Hz (**Fig. 4A**). We found no differences between genotypes in resting membrane potential (RMP, **Fig. 4B**) or action potential amplitude (APA, **Fig. 4C**). However, AP duration (APD) at 20% (APD_20_, **Fig. 4D**), 50% (APD_50_, **Fig. 4E**), and 90% (APD_90_, **Fig. 4F**) repolarization was significantly prolonged in *Mib1^flox^;Tnnt2^Cre^* cardiomyocytes.

**Figure 3.**
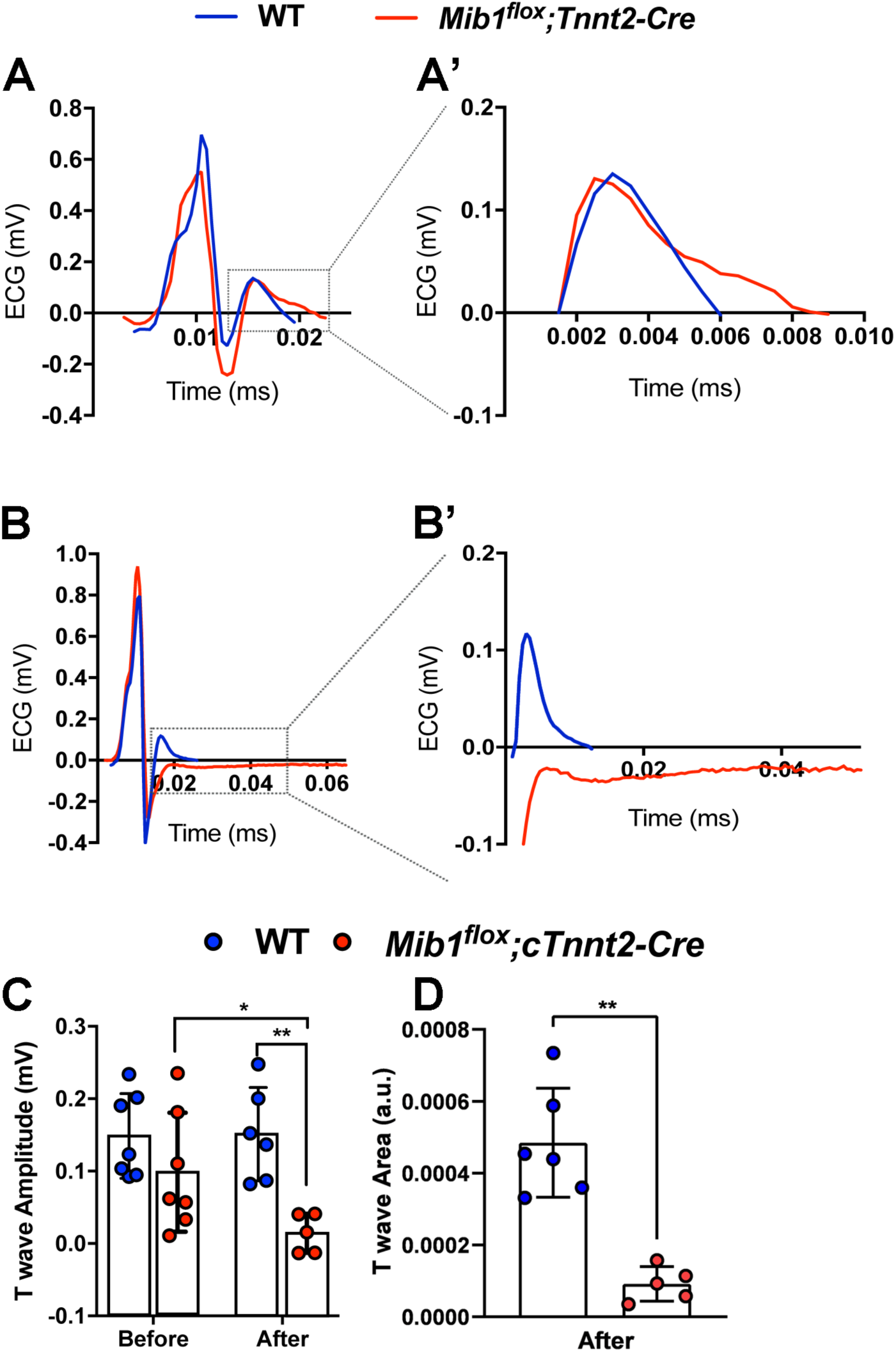
Swimming endurance training reduces T-wave amplitude and area in *Mib1^flox^;Tnnt2^Cre^* mice. (A) Representative QRS complex and T-wave, with (A’) a close-up of the T-wave in WT and *Mib1^flox^;Tnnt2^Cre^* mice before the swimming protocol. (B) Representative QRS complex and T-wave, with (B’) a close-up of the T-wave after the swimming protocol, showing differences in T-wave morphology. (C and D) Summary data showing decreased T-wave amplitude (C) and area (D) in *Mib1^flox^;Tnnt2^Cre^* mice after training. Statistical significance was determined by unpaired two-tailed Student’s t-test.*\*P* < 0.05 Results represent mean±SD of 7-5 animals per genotype.

**Figure 4.**
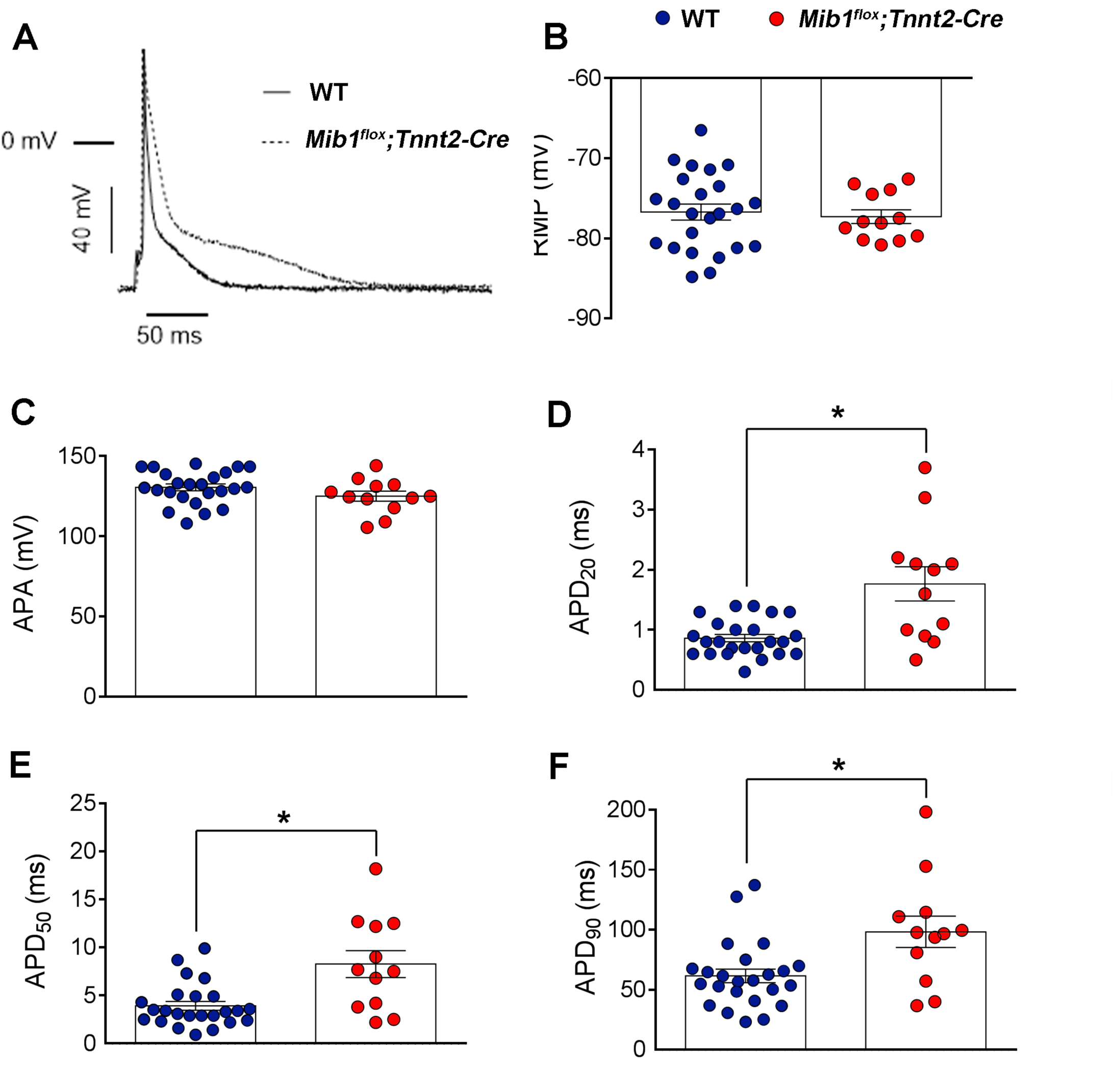
*Mib1* inactivation increases action potential duration. Panels A-F show electrophysiological parameters in cardiomyocytes from WT (n=5) and *Mib1^flox^;Tnnt2^Cre^* mice (n=5). (A) Superimposed action potentials (APs), (B) resting membrane potential (RMP), (C) Action potential amplitude (APA), and (D-F) action potential duration (APD) measured at 20%, 50%, and 90% repolarization. Significant differences (*\*P<0.05* vs WT) in D-F were determined using unpaired two-tailed Student’s t-test, with non-parametric tests (two-sided Wilcoxon’s test) used for small sample sizes (n<15). Data were also analyzed using multilevel mixed-effects models to account for repeated sample assessments. Each bar represents the mean±SEM of 12 and 24 cardiomyocytes from five mice per genotype, with individual data point representing separate experiments.

### Altered cell capacitance and increased peak I_Na_ density in *Mib1^flox^;Tnnt2^Cre^*mice

To assess changes in ionic currents underlying APs, we used patch-clamp recording techniques. Cardiomyocytes from *Mib1^flox^;Tnnt2^Cre^* mice exhibited significantly larger cell capacitance compared to WT (**S3C Fig.**), confirmed by direct measurement of cardiomyocyte area using wheat germ agglutinin staining (**S3A,B Fig.**). We then recorded sodium currents (I_Na_) in isolated cardiomyocytes from both genotypes. **Fig. 5A** and **B** show I_Na_ traces and current-density voltage curve, respectively, revealing that *Mib1^flox^;Tnnt2^Cre^* cardiomyocytes had significantly higher peak I_Na_ density between –50 and –30 mV (**Fig. 5A,B**), with no changes in current activation or inactivation kinetics (**S4 Table**). The voltage dependence of peak I_Na_ activation and inactivation was similar in WT and *Mib1^flox^;Tnnt2^Cre^* cardiomyocytes (**Fig. 5C,D**), with no significant differences in midpoint or slope values of conductance-voltage and inactivation curves (**S4 Table**). **S4 Fig.** shows the time course of peak I_Na_ reactivation, which overlapped between both groups, indicating no change in reactivation kinetics in *Mib1^flox^;Tnnt2^Cre^*cardiomyocytes (**S4 Table**). Additionally, late sodium current (I_NaL_) measured by 500 ms pulses from –120 to –40 mV was also similar in WT and *Mib1^flox^;Tnnt2^Cre^* cardiomyocytes (**S4B Fig.**). Next, we examined potential differences in L-type calcium currents (I_CaL_). I_CaL_ traces were recorded following a protocol shown in **Fig. 5E**, revealing no difference in I_CaL_ density between WT and *Mib1^flox^;Tnnt2^Cre^* cardiomyocytes across all tested potentials (**Fig. 5E,F**). Activation and inactivation kinetics of I_CaL_ were also similar between genotypes (**Fig. 5G, S4 Table**), although inactivation midpoint shifted to more depolarized potentials in *Mib1^flox^;Tnnt2^Cre^* cardiomyocytes (**Fig. 5H**, **Suppl. Table 4**).

**Figure 5.**
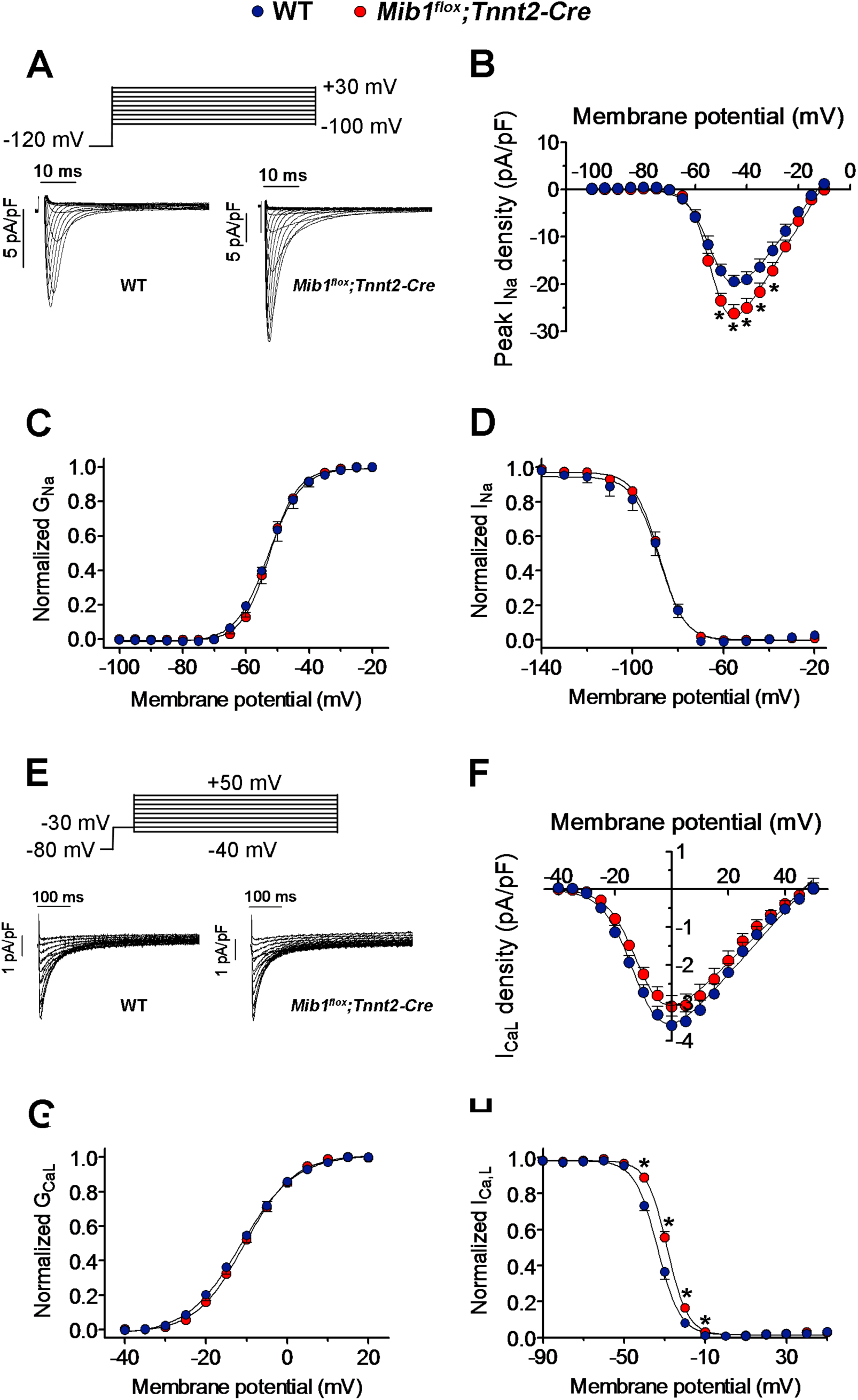
Effects of *Mib1* inactivation on sodium. **(I**_Na_**) and calcium (I**_CaL_**) currents.** (A) I_Na_ traces from cardiomyocytes of WT and *Mib1^flox^;Tnnt2^Cre^* mice, with (B) mean peak I_Na_ density-voltage relationships shown (*\*P<0.05* vs WT). (C,D) Conductance-voltage and inactivation curves for I_Na_, respectively, fitted with Boltzmann functions. Similar analyses for I_CaL_ are shown in panels (E) to (H), with (F) depicting mean I_CaL_ density-voltage relationships. Statistical analyses utilized unpaired two-tailed Student’s t-tests and, for smaller sample sizes, non-parametric tests (two-sided Wilcoxon’s test). Multilevel mixed-effects models were applied for repeated sample assessments. Each point (B-D and F-H) represents the mean±SEM of 17-13 WT and 9 *Mib1^flox^;Tnnt2^Cre^* cardiomyocytes from five mice per genotype.

### Decreased peak and sustained outward K^+^ currents in *Mib1^flox^;Tnnt2^Cre^* mice

In mouse cardiomyocytes, several voltage-dependent K^+^ channels contribute to outward repolarizing currents, including Kv4.3/4.2 and Kv1.5, which are also present in the human myocardium. Kv1.5 is mainly expressed in the atria [35]. We compared K^+^ currents in WT and *Mib Mib1^flox^;Tnnt2^Cre^* cardiomyocytes using 500-ms pulses from –80 and +50 mV (**Fig. 6A**). In *Mib1^flox^;Tnnt2^Cre^* cardiomyocytes showed smaller peak transient and sustained currents compared to WT. Density-voltage curves revealed significant reductions in peak current density at potentials ≥−10 mV (**Fig. 6B)** and sustained current at potentials ≥−20 mV (**Fig. 6C**), with a 45.4% reduction in sustained current density +50 mV, and a 25% reduction in peak current density at this voltage (*P*<0.05). The decay of outward K^+^ current was faster in *Mib1^flox^;Tnnt2^Cre^* cardiomyocytes (**Fig. 6A**, inset), with a significantly reduced time constant (**Fig. 6D**, **S4 Table**). To further characterize outward K^+^ currents in *Mib1^flox^;Tnnt2^Cre^* cardiomyocytes, we measured the charge (the total amount of K^+^ crossing the membrane) during the 500-ms pulses by integrating the current traces to calculate the area. The charge density at potentials ≥-20 mV was significantly lower in *Mib1^flox^;Tnnt2^Cre^* cardiomyocytes (**Fig. 6E**). However, the inward rectifier current (I_K1_) density and kinetics were similar between WT and *Mib1^flox^;Tnnt2^Cre^* cardiomyocytes (**S5 Fig.**).

**Figure 6.**
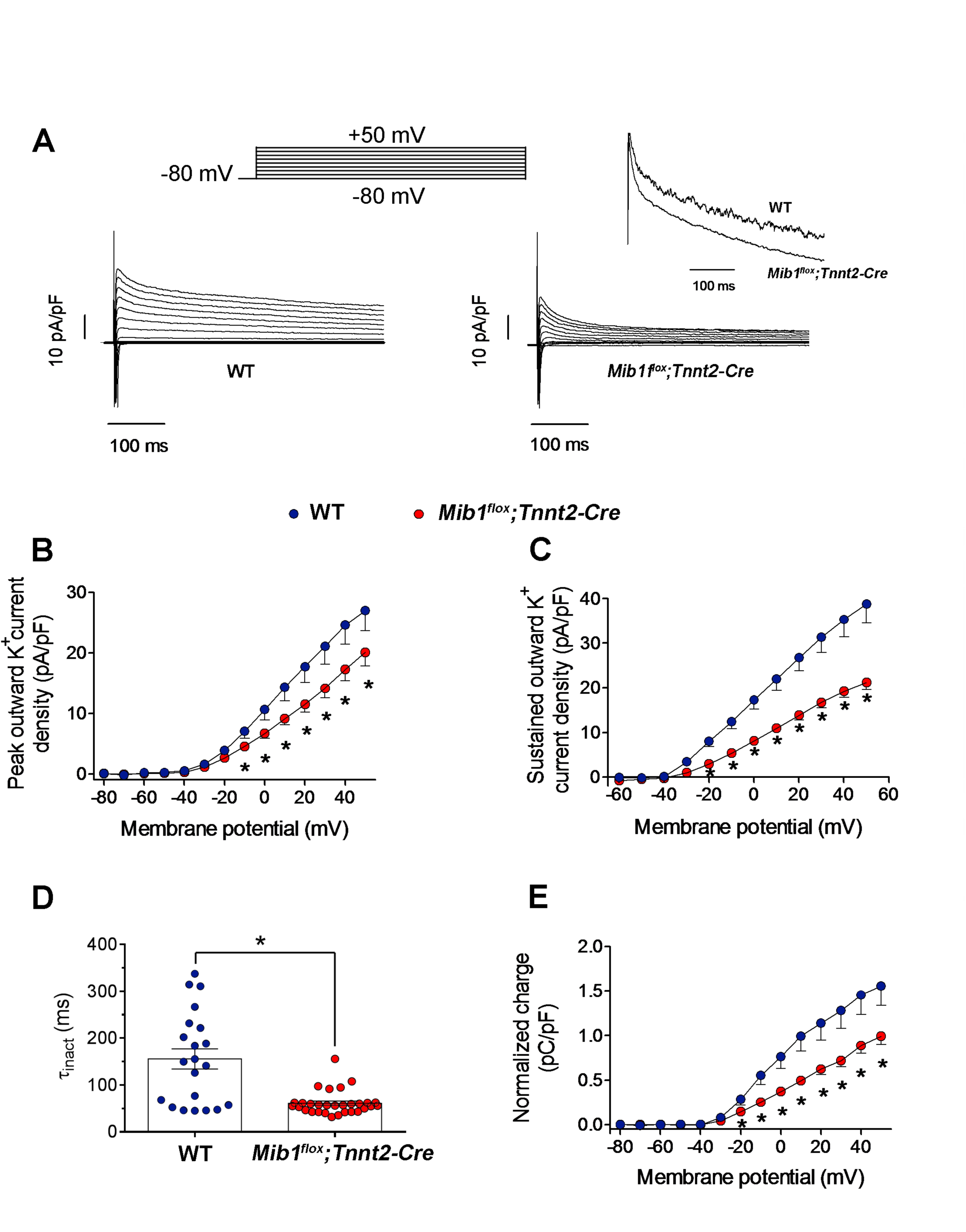
Effects of *Mib1* inactivation on outward K^+^ currents. (A) Outward K^+^ current traces from WT and *Mib1^flox^;Tnnt2^Cre^* cardiomyocytes, with inset highlighting normalized current traces at +50 mV. (B and C) Current density-voltage relationships at peak (B) and end (C) of pulses in both genotypes. (D) Time constants (τ_inact_) of current decay at +50 mV. Each point represents an experiment. (E) Mean outward K^+^ charge density-voltage relationships. Statistical significance (*\*P<0.05* vs WT) was determined using unpaired two-tailed Student’s t-test, adjusting for repeated measures with multilevel mixed-effects models. Data in panels B to E, represent mean±SEM of 21 WT and 30 *Mib1^flox^;Tnnt2^Cre^* cardiomyocytes from five mice per genotype.

### Impaired ion channel expression in *Mib1^flox^;Tnnt2^Cre^* mice

To determine whether the electrophysiological abnormalities in *Mib1^flox^;Tnnt2^Cre^* mice were due to defective ion channel expression, we performed immunofluorescence staining for Na_V_1.5, K_V_4.2, and K_V_4.3 in freshly isolated cardiomyocytes (**Fig. 7**). Na_V_1.5 expression showed no difference between genotypes in integrated density (**Fig. 7A,B**), but protein levels were increased (**Fig.7C,D**). *Mib1^flox^;Tnnt2-Cre* cardiomyocytes had lower integrated densities for K_V_4.2 (**Fig. 7E,F**) and K_V_4.3 (**Fig. 7I,J**), though protein were not significantly different from WT (**Fig. 7G,H,K,L**). These decreases in K_V_4.2 and K_V_4.3 density likely contribute to the reduced outward K^+^ currents and increased APD in *Mib1^flox^;Tnnt2^Cre^* mice.

**Figure 7.**
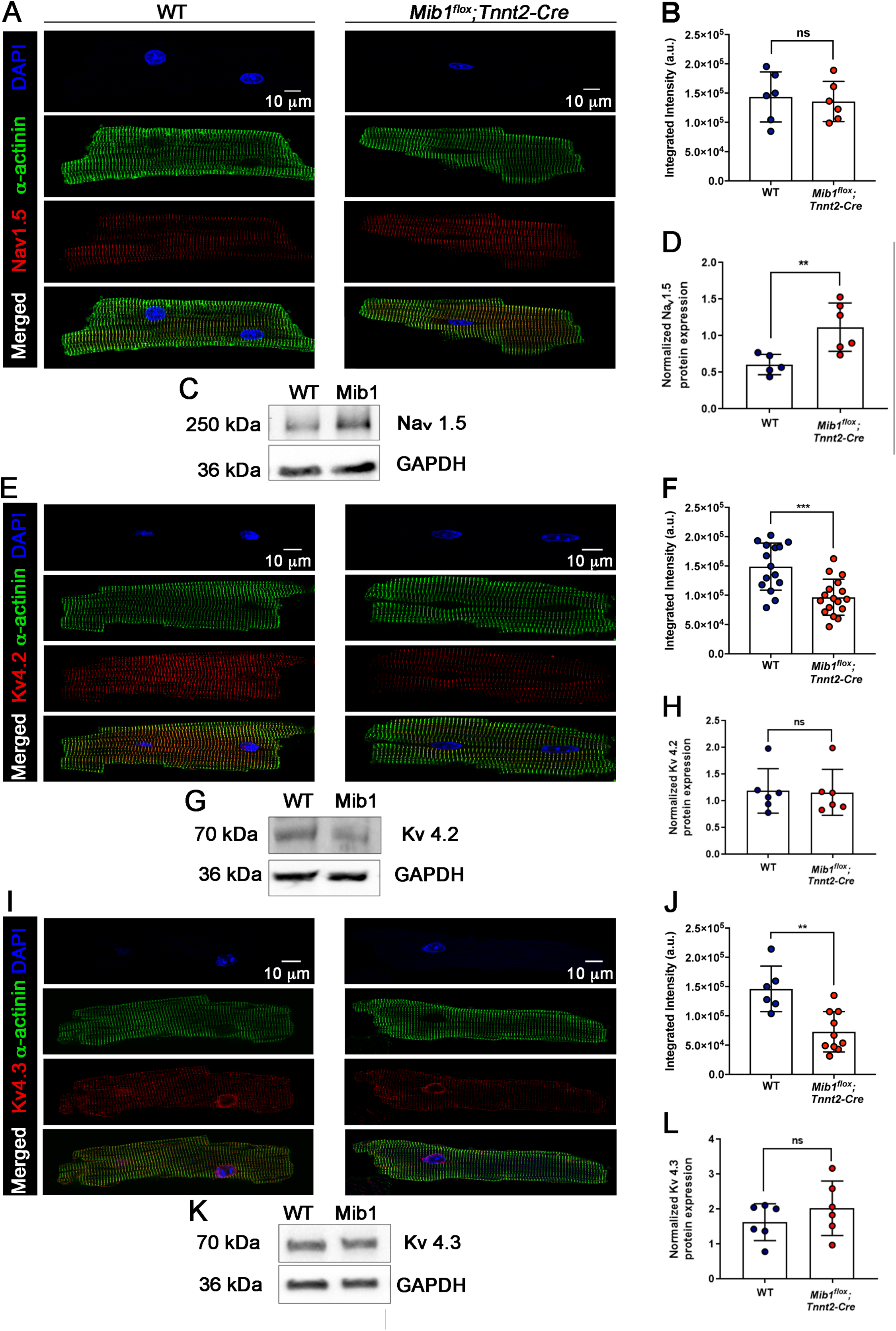
Confocal immunofluorescence images of Nav1.5, Kv4.2, and Kv4.3 localization in isolated cardiomyocytes. (A) Immunofluorescence staining of α-actinin (green) and Nav1.5 (red) in WT and *Mib1^flox^;Tnnt2^Cre^* mice, with merged images showing colocalization. Nuclei were stained with DAPI (blue). (B) Quantification of integrated fluorescence intensity (a.u.) from 6 WT and 6 *Mib1^flox^;Tnnt2^Cre^* cardiomyocytes per genotype. (C) Staining for α-actinin (green) and Kv4.2 (red) with merged images indicating colocalization. (D) Quantification of fluorescence intensity from 15 WT and 17 *Mib1^flox^;Tnnt2^Cre^* cardiomyocytes, showing reduced intensity in *Mib1^flox^;Tnnt2^Cre^* mice (*\*\*\*P* < 0.001 vs WT). (E) Staining for α-actinin (green) and Kv4.3 (red), with merged images highlighting colocalization. (F) Quantification of fluorescence intensity from 6 WT and 10 *Mib1^flox^;Tnnt2^Cre^* cardiomyocytes, showing reduced intensity in *Mib1^flox^;Tnnt2^Cre^* mice (*\*\*P* < 0.01 vs WT). Statistical significance was determined using unpaired two-tailed Student’s t-test. Data represent mean±SD of cells from 3-4 animals per genotype.

### Abnormal Connexin-43 localization in the hearts of *Mib1^flox^;Tnnt2^Cre^* mice

The gap-junction protein Connexin-43 (CX43) is crucial for electrical excitation in cardiomyocytes [36] and requires N-Cadherin for proper localization at intercalated disks [37]. N-Cadherin and Cx43 immunofluorescence analysis (**Fig. 8A-D**) showed significant mislocalization of CX43 to the lateral sides of cardiomyocytes in *Mib1^flox^;Tnnt2^Cre^* hearts (**Fig. 8E-F’**) with reduced CX43 staining associated with N-cadherin (**Fig. 8AG**). Cardiac fibrosis, a potential cause of electrical abnormalities[38], was assessed with Picrosirius red staining, but no significant differences were found between groups, either at baseline or after endurance training and isoproterenol treatment (**S6 Fig. 6**).

**Figure 8.**
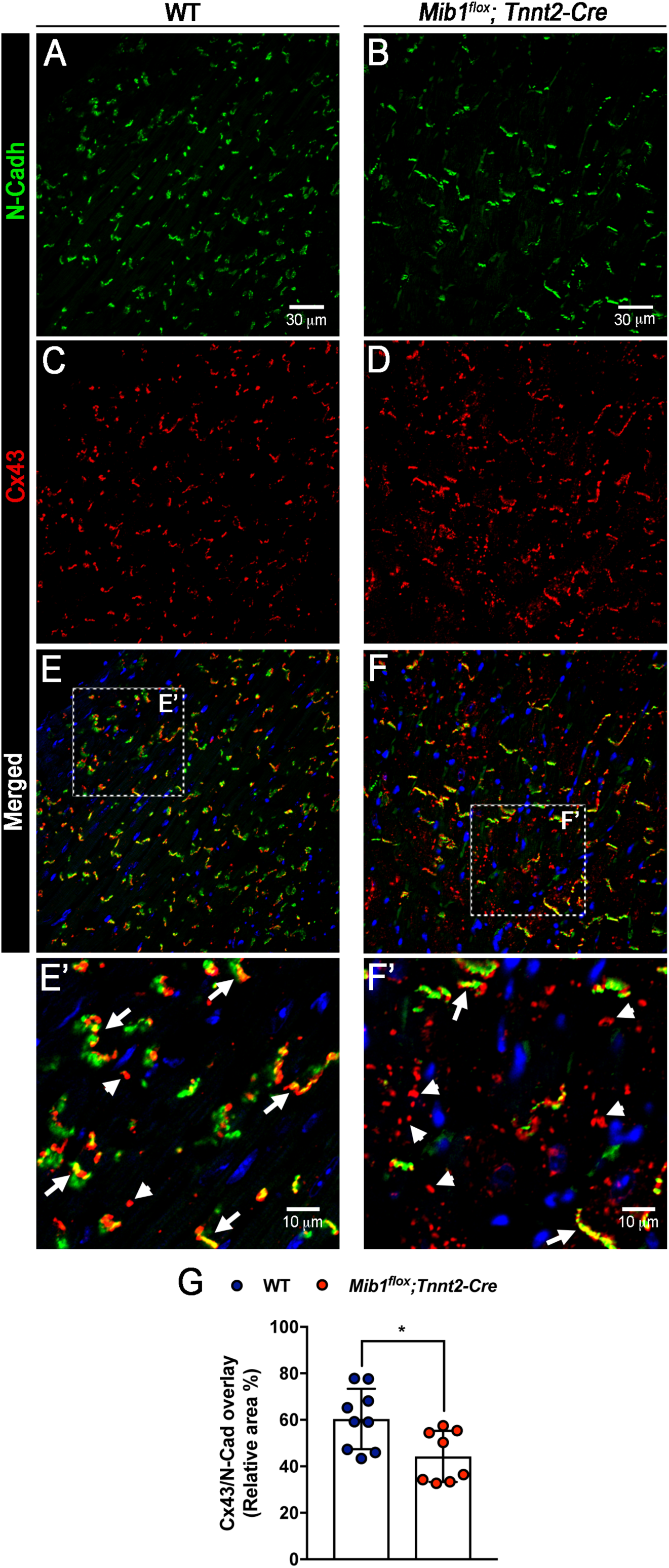
N-cadherin (N-Cadh) and connexin 43 (CX43) expression patterns in left ventricle cardiomyocytes. (A, B) N-Cadh expression (green) at intercalated disks in WT and *Mib1^flox^;Tnnt2^Cre^* cardiomyocytes. (C) CX43 localization (red) at intercalated disks and lateral membranes in WT cardiomyocytes. (D) CX43 expression is concentrated at the lateral membrane in *Mib1^flox^;Tnnt2^Cre^* cardiomyocytes. (E,F) Merged images with DAPI-stained nuclei (blue); arrowheads highlight predominant CX43 expression in WT (E’) and *Mib1^flox^;Tnnt2^Cre^* (F’) cardiomyocytes. (G) Quantification of CX43/N-Cadh overlay at intercalated disks using automated image segmentation. Statistical significance (\**P < 0.05* vs WT) was determined using unpaired two-tailed Student’s t-test. Data represent mean±SD of sections from three animals per genotype, totaling 8-9 sections.

## Discussion

MIB1 is an E3 ubiquitin ligase regulating NOTCH ligands endocytosis[23]. Conditional *Mib1* inactivation in mouse hearts produces an LVNC phenotype[22]. Our study investigated electrical defects in adult mice with *Mib1* inactivation, revealing significant cardiac electrical abnormalities and related molecular, cellular, tissue, and organ changes. *Mib1^flox^;Tnnt2^Cre^* cardiomyocytes showed reduced Kv4.2 and Kv4.3 ion channel expression, increased peak I_Na_ density, reduced outward K^+^ currents densities, and prolonged APD. Under cardiac stress from isoproterenol or swimming, these mice exhibited prolonged QTc duration and decreased T-wave amplitude and area. Patients with MIB1 mutations showed similar prolonged QTc duration and increased QT variability.

LVNC often features abnormal ECG patterns, such as intraventricular conduction delays, left ventricular hypertrophy, left axis deviation, repolarization abnormalities, and QTc prolongation [27]. In mice with LVNC resulting from Nkx2-5 inactivation, defects include a widened QRS interval and reduced R amplitude, but no changes in QT interval or T wave amplitude [13]. In our study, *Mib1^flox^;Tnnt2^Cre^* mice showed no significant changes in basal cardiac electrical activity but tended towards a lengthened PR interval and QTc duration. ECG abnormalities can appear during exercise or with symptoms; one LVNC patient had exercise-induced atrial and ventricular tachycardia [39]. Swimming endurance caused a ∼2 ms increase in QRS interval and ∼8.8 ms increase in QTc_90_ duration in *Mib1^flox^;Tnnt2^Cre^* mice. Isoproterenol-induced stress increased QTc_90_ duration by ∼5.8 ms. Prolonged QT interval raises the risk of ventricular arrhythmia and sudden cardiac death [40]. Four *Mib1^flox^;Tnnt2^Cre^* mice died during the swimming protocol, likely due to ventricular repolarization abnormalities. *Mib1^flox^;Tnnt2^Cre^* mutants have reduced ejection fraction and impaired coronary vessel formation [22], triggering ventricular ischemia. Hypoxia in adult patients is linked to prolonged QT interval and arrhythmias [41]. Neonatal mice with oxygen deprivation or increased hypoxia-inducible factor signaling showed prolonged QTc, increased QTc dispersion, disturbed ion channel expression, and higher sudden death risk [42]. During exercise, hypoxia is associated with lower P-wave, QRS complex, and T-wave amplitudes [43].

The T-wave in the ECG indicates ventricular repolarization, with alterations like increased duration, low amplitude, and inversion. In *Mib1^flox^;Tnnt2^Cre^* mice, post-swimming T-wave analysis showed low amplitude and area, indicating ventricular repolarization abnormalities. These may result from low K_V_4.2 and K_V_4.3 ion channel density, reduced outward K^+^ current, and increased AP duration. Low transient outward K^+^ current and Kv4.3 mRNA expression are linked to low T-wave amplitude in long-term cardiac memory models [44].

Cardiac APD is regulated by depolarizing currents (mainly I_Na_ and I_Ca,L_) and hyperpolarizing currents (transient outward K^+^, delayed rectifier, and I_K1_ currents) [45]. I_Na_ controls rapid depolarization of cardiomyocytes, and I_NaL_ regulates action potential duration[46]. Our results showed a significant increase in peak I_Na_ and Nav1.5 expression in *Mib1^flox^;Tnnt2^Cre^* cardiomyocytes. Persistent PKA activation promotes Nav1.5 trafficking to the cytoplasmic membrane, especially at intercalated discs [47]. This increase in I_Na_ could be due to persistent β-adrenoceptor stimulation from cardiac stress. However, this did not enhance intracardiac conduction; instead, the QRS interval was widened in both LVNC patients and *Mib1^flox^;Tnnt2^Cre^* mice post-swimming.

Gap junction proteins, essential for intercellular electrical coupling, are mainly located at intercalated disks[48]. CX43 is the primary connexin in ventricles, and its impaired function is linked to many cardiomyopathies [49]. In *Mib1^flox^;Tnnt2^Cre^* cardiomyocytes, CX43 was mostly located laterally, unlike its typical position at intercalated discs in WT cells. This abnormal distribution likely contributes to conduction defects [50], overshadowing the increased INa and explaining the conduction defects in *Mib1^flox^;Tnnt2^Cre^* mice and LVNC patients.

Studies on ICa,L in heart failure have reported conflicting results[51, 52]. After normalizing for cell capacitance, we found no significant differences in peak ICa,L density or inactivation kinetics between genotypes. However, in *Mib1^flox^;Tnnt2^Cre^* cardiomyocytes, the inactivation-curve midpoint was slightly shifted to more depolarized potentials. Thus, the prolonged APD in *Mib1^flox^;Tnnt2^Cre^* cardiomyocytes is likely due to changes in currents other than L-type Ca2+ channels.

A decrease in outward K^+^ current prolongs the APD and QT interval [53]. *Mib1^flox^;Tnnt2^Cre^* mice had significantly reduced peak and sustained outward K^+^ currents, closely associated with increased APD_20_, APD_50_, and APD_90_. Kv4.3/4.2 channel expression was significantly reduced, leading to lower transient outward K^+^ current, increased Ca^2+^ influx, and higher intracellular calcium concentration, enhancing CaMKII activity[35]. This activates CaMKII and calcineurin, triggering hypertrophic gene activation as a compensatory response, but sustained hypertrophy risks progressing to heart failure and cardiomyopathy[54].

QT variability measures spontaneous fluctuations in QT interval duration [55] and is an indicator of ventricular arrhythmias and sudden cardiac death. Increased QT variability is linked to coronary artery disease and left ventricular hypertrophy [56]. While QT variability was not increased in *Mib1^flox^;Tnnt2^Cre^* mice, it was elevated in LVNC patients with *MIB1* mutations, suggesting a higher risk of ventricular arrhythmias. Other features of *Mib1^flox^;Tnnt2^Cre^* mice, such as a long QTc, were also present in LVNC patients with *MIB1* mutations.

## Conclusions

*Mib1^flox^;Tnnt2^Cre^* mice exhibit prolonged QTc duration and decreased T-wave amplitude and area under swimming endurance exercise or isoproterenol-induced stress. Freshly isolated *Mib1^flox^;Tnnt2^Cre^* cardiomyocytes showed increased I_Na_, decreased outward K^+^ currents, prolonged APD, reduced Kv4.2 and KV4.3 expression, and lateral redistribution of CX43. LVNC patients with MIB1 mutations also show prolonged QTc interval and increased QT variability markers.

## References

1. Oechslin E, Jenni R. Left ventricular non-compaction revisited: a distinct phenotype with genetic heterogeneity? Eur Heart J. 2011;32(12):1446–56. Epub 2011/02/03. doi: ehq508 [pii] 10.1093/eurheartj/ehq508. PubMed PMID: 21285074.

2. Gati S, Papadakis M, Papamichael ND, Zaidi A, Sheikh N, Reed M, et al. Reversible de novo left ventricular trabeculations in pregnant women: implications for the diagnosis of left ventricular noncompaction in low-risk populations. Circulation. 2014;130(6):475–83. Epub 20140708. doi: 10.1161/CIRCULATIONAHA.114.008554. PubMed PMID: 25006201.

3. Wengrofsky P, Armenia C, Oleszak F, Kupferstein E, Rednam C, Mitre CA, et al. Left Ventricular Trabeculation and Noncompaction Cardiomyopathy: A Review. EC Clin Exp Anat. 2019;2(6):267–83. Epub 20190729. PubMed PMID: 31799511; PubMed Central PMCID: PMCPMC6890222.

4. Duru F, Candinas R. Noncompaction of ventricular myocardium and arrhythmias. J Cardiovasc Electrophysiol. 2000;11(4):493. doi: 10.1111/j.1540-8167.2000.tb00350.x. PubMed PMID: 10809508.

5. Ichida F. Left ventricular noncompaction – Risk stratification and genetic consideration. J Cardiol. 2020;75(1):1–9. Epub 20191017. doi: 10.1016/j.jjcc.2019.09.011. PubMed PMID: 31629663.

6. Pignatelli RH, McMahon CJ, Dreyer WJ, Denfield SW, Price J, Belmont JW, et al. Clinical characterization of left ventricular noncompaction in children: a relatively common form of cardiomyopathy. Circulation. 2003;108(21):2672–8. Epub 2003/11/19. doi: 10.1161/01.CIR.0000100664.10777.B8. PubMed PMID: 14623814.

7. Klaassen S, Probst S, Oechslin E, Gerull B, Krings G, Schuler P, et al. Mutations in sarcomere protein genes in left ventricular noncompaction. Circulation. 2008;117(22):2893–901. Epub 2008/05/29. doi: 10.1161/CIRCULATIONAHA.107.746164. PubMed PMID: 18506004.

8. van Waning JI, Caliskan K, Hoedemaekers YM, van Spaendonck-Zwarts KY, Baas AF, Boekholdt SM, et al. Genetics, Clinical Features, and Long-Term Outcome of Noncompaction Cardiomyopathy. J Am Coll Cardiol. 2018;71(7):711–22. Epub 2018/02/16. doi: 10.1016/j.jacc.2017.12.019. PubMed PMID: 29447731.

9. Ichida F, Tsubata S, Bowles KR, Haneda N, Uese K, Miyawaki T, et al. Novel gene mutations in patients with left ventricular noncompaction or Barth syndrome. Circulation. 2001;103(9):1256–63. Epub 2001/03/10. PubMed PMID: 11238270.

10. Grego-Bessa J, Luna-Zurita L, del Monte G, Bolos V, Melgar P, Arandilla A, et al. Notch signaling is essential for ventricular chamber development. Dev Cell. 2007;12(3):415–29. Epub 2007/03/06. doi: S1534-5807(06)00600-9 [pii] 10.1016/j.devcel.2006.12.011. PubMed PMID: 17336907.

11. Sedaghat-Hamedani F, Haas J, Zhu F, Geier C, Kayvanpour E, Liss M, et al. Clinical genetics and outcome of left ventricular non-compaction cardiomyopathy. Eur Heart J. 2017;38(46):3449–60. doi: 10.1093/eurheartj/ehx545. PubMed PMID: 29029073.

12. Zhang W, Chen H, Qu X, Chang CP, Shou W. Molecular mechanism of ventricular trabeculation/compaction and the pathogenesis of the left ventricular noncompaction cardiomyopathy (LVNC). Am J Med Genet C Semin Med Genet. 2013;163C(3):144-56. Epub 2013/07/12. doi: 10.1002/ajmg.c.31369. PubMed PMID: 23843320; PubMed Central PMCID: PMCPMC3725649.

13. Choquet C, Nguyen THM, Sicard P, Buttigieg E, Tran TT, Kober F, et al. Deletion of Nkx2-5 in trabecular myocardium reveals the developmental origins of pathological heterogeneity associated with ventricular non-compaction cardiomyopathy. PLoS Genet. 2018;14(7):e1007502. Epub 2018/07/07. doi: 10.1371/journal.pgen.1007502. PubMed PMID: 29979676; PubMed Central PMCID: PMCPMC6051668.

14. Artavanis-Tsakonas S, Rand MD, Lake RJ. Notch signaling: cell fate control and signal integration in development. Science. 1999;284(5415):770-6.

15. Kopan R. Notch signaling. Cold Spring Harbor perspectives in biology. 2012;4(10). Epub 2012/10/03. doi: 10.1101/cshperspect.a011213. PubMed PMID: 23028119.

16. Garg V, Muth AN, Ransom JF, Schluterman MK, Barnes R, King IN, et al. Mutations in NOTCH1 cause aortic valve disease. Nature. 2005;437(7056):270–4. PubMed PMID: 16025100.

17. Joutel A, Corpechot C, Ducros A, Vahedi K, Chabriat H, Mouton P, et al. Notch3 mutations in CADASIL, a hereditary adult-onset condition causing stroke and dementia. Nature. 1996;383(6602):707-10. PubMed PMID: 8878478.

18. Li L, Krantz ID, Deng Y, Genin A, Banta AB, Collins CC, et al. Alagille syndrome is caused by mutations in human Jagged1, which encodes a ligand for Notch1. Nat Genet. 1997;16(3):243–51. PubMed PMID: 9207788.

19. Oda T, Elkahloun AG, Pike BL, Okajima K, Krantz ID, Genin A, et al. Mutations in the human Jagged1 gene are responsible for Alagille syndrome. Nat Genet. 1997;16(3):235–42. lineage decision. PubMed PMID: 9207787.

20. MacGrogan D, Munch J, de la Pompa JL. Notch and interacting signalling pathways in cardiac development, disease, and regeneration. Nat Rev Cardiol. 2018;15(11):685–704. Epub 2018/10/06. doi: 10.1038/s41569-018-0100-2. PubMed PMID: 30287945.

21. Meester JAN, Verstraeten A, Alaerts M, Schepers D, Van Laer L, Loeys BL. Overlapping but distinct roles for NOTCH receptors in human cardiovascular disease. Clin Genet. 2019;95(1):85–94. Epub 2018/05/17. doi: 10.1111/cge.13382. PubMed PMID: 29767458.

22. Luxan G, Casanova JC, Martinez-Poveda B, Prados B, D’Amato G, MacGrogan D, et al. Mutations in the NOTCH pathway regulator MIB1 cause left ventricular noncompaction cardiomyopathy. Nature medicine. 2013;19(2):193–201. Epub 2013/01/15. doi: 10.1038/nm.3046. PubMed PMID: 23314057.

23. Itoh M, Kim CH, Palardy G, Oda T, Jiang YJ, Maust D, et al. Mind bomb is a ubiquitin ligase that is essential for efficient activation of Notch signaling by Delta. Dev Cell. 2003;4(1):67–82. PubMed PMID: 12530964.

24. D’Amato G, Luxan G, de la Pompa JL. Notch signalling in ventricular chamber development and cardiomyopathy. FEBS J. 2016;283(23):4223–37. Epub 20160622. doi: 10.1111/febs.13773. PubMed PMID: 27260948.

25. Siguero-Alvarez M, Salguero-Jimenez A, Grego-Bessa J, de la Barrera J, MacGrogan D, Prados B, et al. A Human Hereditary Cardiomyopathy Shares a Genetic Substrate With Bicuspid Aortic Valve. Circulation. 2023;147(1):47–65. Epub 20221103. doi: 10.1161/CIRCULATIONAHA.121.058767. PubMed PMID: 36325906.

26. Steffel J, Kobza R, Oechslin E, Jenni R, Duru F. Electrocardiographic characteristics at initial diagnosis in patients with isolated left ventricular noncompaction. Am J Cardiol. 2009;104(7):984–9. doi: 10.1016/j.amjcard.2009.05.042. PubMed PMID: 19766768.

27. Towbin JA, Lorts A, Jefferies JL. Left ventricular non-compaction cardiomyopathy. Lancet. 2015;386(9995):813-25. doi: 10.1016/S0140-6736(14)61282-4. PubMed PMID: 25865865.

28. Rivera-Torres J, Calvo CJ, Llach A, Guzman-Martinez G, Caballero R, Gonzalez-Gomez C, et al. Cardiac electrical defects in progeroid mice and Hutchinson-Gilford progeria syndrome patients with nuclear lamina alterations. Proc Natl Acad Sci U S A. 2016;113(46):E7250–E9. Epub 20161031. doi: 10.1073/pnas.1603754113. PubMed PMID: 27799555; PubMed Central PMCID: PMCPMC5135377.

29. Mitchell GF, Jeron A, Koren G. Measurement of heart rate and Q-T interval in the conscious mouse. Am J Physiol. 1998;274(3):H747–51. doi: 10.1152/ajpheart.1998.274.3.H747. PubMed PMID: 9530184.

30. Cruz FM, Sanz-Rosa D, Roche-Molina M, Garcia-Prieto J, Garcia-Ruiz JM, Pizarro G, et al. Exercise triggers ARVC phenotype in mice expressing a disease-causing mutated version of human plakophilin-2. J Am Coll Cardiol. 2015;65(14):1438–50. doi: 10.1016/j.jacc.2015.01.045. PubMed PMID: 25857910.

31. Puhl SL, Weeks KL, Ranieri A, Avkiran M. Assessing structural and functional responses of murine hearts to acute and sustained beta-adrenergic stimulation in vivo. J Pharmacol Toxicol Methods. 2016;79:60–71. Epub 20160204. doi: 10.1016/j.vascn.2016.01.007. PubMed PMID: 26836145; PubMed Central PMCID: PMCPMC4840275.

32. Nieto-Marín P, Tinaquero D, Utrilla RG, Cebrián J, González-Guerra A, Crespo-García T, et al. Tbx5 variants disrupt Nav1.5 function differently in patients diagnosed with Brugada or Long QT Syndrome. Cardiovasc Res. 2022;118(4):1046–60. doi: 10.1093/cvr/cvab045. PubMed PMID: 33576403.

33. Caballero R, Utrilla RG, Amoros I, Matamoros M, Perez-Hernandez M, Tinaquero D, et al. Tbx20 controls the expression of the KCNH2 gene and of hERG channels. Proc Natl Acad Sci U S A. 2017;114(3):E416–E25. Epub 20170103. doi: 10.1073/pnas.1612383114. PubMed PMID: 28049825; PubMed Central PMCID: PMCPMC5255604.

34. Le Page S, Niro M, Fauconnier J, Cellier L, Tamareille S, Gharib A, et al. Increase in Cardiac Ischemia-Reperfusion Injuries in Opa1+/− Mouse Model. PLoS One. 2016;11(10):e0164066. Epub 20161010. doi: 10.1371/journal.pone.0164066. PubMed PMID: 27723783; PubMed Central PMCID: PMCPMC5056696.

35. Nerbonne JM, Kass RS. Molecular physiology of cardiac repolarization. Physiol Rev. 2005;85(4):1205–53. doi: 10.1152/physrev.00002.2005. PubMed PMID: 16183911.

36. Beauchamp P, Desplantez T, McCain ML, Li W, Asimaki A, Rigoli G, et al. Electrical coupling and propagation in engineered ventricular myocardium with heterogeneous expression of connexin43. Circ Res. 2012;110(11):1445–53. Epub 20120419. doi: 10.1161/CIRCRESAHA.111.259705. PubMed PMID: 22518032; PubMed Central PMCID: PMCPMC3381798.

37. Matsuda T, Fujio Y, Nariai T, Ito T, Yamane M, Takatani T, et al. N-cadherin signals through Rac1 determine the localization of connexin 43 in cardiac myocytes. J Mol Cell Cardiol. 2006;40(4):495–502. Epub 20060302. doi: 10.1016/j.yjmcc.2005.12.010. PubMed PMID: 16515795.

38. Nguyen TP, Qu Z, Weiss JN. Cardiac fibrosis and arrhythmogenesis: the road to repair is paved with perils. J Mol Cell Cardiol. 2014;70:83–91. Epub 20131031. doi: 10.1016/j.yjmcc.2013.10.018. PubMed PMID: 24184999; PubMed Central PMCID: PMCPMC3995831.

39. Seethala S, Knollman F, McNamara D, Saba S, Schwender F, Schwartzman D, et al. Exercise-induced atrial and ventricular tachycardias in a patient with left ventricular noncompaction and normal ejection fraction. Pacing Clin Electrophysiol. 2011;34(10):e94–7. Epub 20100621. doi: 10.1111/j.1540-8159.2010.02813.x. PubMed PMID: 20565692.

40. Postema PG, Wilde AA. The measurement of the QT interval. Curr Cardiol Rev. 2014;10(3):287–94. doi: 10.2174/1573403x10666140514103612. PubMed PMID: 24827793; PubMed Central PMCID: PMCPMC4040880.

41. Roche F, Reynaud C, Pichot V, Duverney D, Costes F, Garet M, et al. Effect of acute hypoxia on QT rate dependence and corrected QT interval in healthy subjects. Am J Cardiol. 2003;91(7):916–9. doi: 10.1016/s0002-9149(03)00040-7. PubMed PMID: 12667592.

42. Neary MT, Mohun TJ, Breckenridge RA. A mouse model to study the link between hypoxia, long QT interval and sudden infant death syndrome. Dis Model Mech. 2013;6(2):503–7. Epub 20120911. doi: 10.1242/dmm.010587. PubMed PMID: 22977222; PubMed Central PMCID: PMCPMC3597031.

43. Coustet B, Lhuissier FJ, Vincent R, Richalet JP. Electrocardiographic changes during exercise in acute hypoxia and susceptibility to severe high-altitude illnesses. Circulation. 2015;131(9):786–94. Epub 20150105. doi: 10.1161/CIRCULATIONAHA.114.013144. PubMed PMID: 25561515.

44. Yu H, McKinnon D, Dixon JE, Gao J, Wymore R, Cohen IS, et al. Transient outward current, Ito1, is altered in cardiac memory. Circulation. 1999;99(14):1898–905. doi: 10.1161/01.cir.99.14.1898. PubMed PMID: 10199889.

45. Wickenden AD, Kaprielian R, Kassiri Z, Tsoporis JN, Tsushima R, Fishman GI, et al. The role of action potential prolongation and altered intracellular calcium handling in the pathogenesis of heart failure. Cardiovasc Res. 1998;37(2):312–23. doi: 10.1016/s0008-6363(97)00256-3. PubMed PMID: 9614488.

46. Ke HY, Yang HY, Francis AJ, Collins TP, Surendran H, Alvarez-Laviada A, et al. Changes in cellular Ca(2+) and Na(+) regulation during the progression towards heart failure in the guinea pig. J Physiol. 2020;598(7):1339–59. Epub 20190318. doi: 10.1113/JP277038. PubMed PMID: 30811606; PubMed Central PMCID: PMCPMC7187457.

47. Bernas T, Seo, J., Wilson, Z. T., Tan, B. H., Deschenes, I., Carter, C., Liu, J., Tseng, G. N. Persistent PKA activation redistributes NaV1.5 to the cell surface of adult rat ventricular myocytes J Gen Physiol. 2024;156(2):e202313436. doi: 10.1085/jgp.202313436.

48. Vaidya D, Tamaddon HS, Lo CW, Taffet SM, Delmar M, Morley GE, et al. Null mutation of connexin43 causes slow propagation of ventricular activation in the late stages of mouse embryonic development. Circ Res. 2001;88(11):1196–202. doi: 10.1161/hh1101.091107. PubMed PMID: 11397787.

49. Seidel T, Salameh A, Dhein S. A simulation study of cellular hypertrophy and connexin lateralization in cardiac tissue. Biophys J. 2010;99(9):2821–30. doi: 10.1016/j.bpj.2010.09.010. PubMed PMID: 21044579; PubMed Central PMCID: PMCPMC2965950.

50. Gutstein DE, Morley GE, Tamaddon H, Vaidya D, Schneider MD, Chen J, et al. Conduction slowing and sudden arrhythmic death in mice with cardiac-restricted inactivation of connexin43. Circ Res. 2001;88(3):333–9. doi: 10.1161/01.res.88.3.333. PubMed PMID: 11179202; PubMed Central PMCID: PMCPMC3630465.

51. Beuckelmann DJ, Nabauer M, Erdmann E. Intracellular calcium handling in isolated ventricular myocytes from patients with terminal heart failure. Circulation. 1992;85(3):1046–55. doi: 10.1161/01.cir.85.3.1046. PubMed PMID: 1311223.

52. Qin D, Zhang ZH, Caref EB, Boutjdir M, Jain P, el-Sherif N. Cellular and ionic basis of arrhythmias in postinfarction remodeled ventricular myocardium. Circ Res. 1996;79(3):461–73. doi: 10.1161/01.res.79.3.461. PubMed PMID: 8781480.

53. Qu Z, Chung D. Mechanisms and determinants of ultralong action potential duration and slow rate-dependence in cardiac myocytes. PLoS One. 2012;7(8):e43587. Epub 20120827. doi: 10.1371/journal.pone.0043587. PubMed PMID: 22952713; PubMed Central PMCID: PMCPMC3428352.

54. Molkentin JD, Lu JR, Antos CL, Markham B, Richardson J, Robbins J, et al. A calcineurin-dependent transcriptional pathway for cardiac hypertrophy. Cell. 1998;93(2):215–28. PubMed PMID: 9568714.

55. Berger RD, Kasper EK, Baughman KL, Marban E, Calkins H, Tomaselli GF. Beat-to-beat QT interval variability: novel evidence for repolarization lability in ischemic and nonischemic dilated cardiomyopathy. Circulation. 1997;96(5):1557–65. doi: 10.1161/01.cir.96.5.1557. PubMed PMID: 9315547.

56. Schlegel TT, Kulecz WB, Feiveson AH, Greco EC, DePalma JL, Starc V, et al. Accuracy of advanced versus strictly conventional 12-lead ECG for detection and screening of coronary artery disease, left ventricular hypertrophy and left ventricular systolic dysfunction. BMC Cardiovasc Disord. 2010;10:28. Epub 20100616. doi: 10.1186/1471-2261-10-28. PubMed PMID: 20565702; PubMed Central PMCID: PMCPMC2894002.

