## Supplemental file for "Cardiac electrical abnormalities in a mouse model of left ventricular non-compaction cardiomyopathy"

**Short Title: Cardiac electrical abnormalities in LVNC**

Vítor S. Fernandes, PhD[1,2], Ricardo Caballero, PhD[2,3], Marcos Siguero-Álvarez, PhD[1,2], Tania Papoutsi[1], PhD, Juan Ramón Gimeno, MD, PhD[2,4], Eva Delpón, PhD[2,3] & José Luís de la Pompa, PhD[1,2]

[1]Intercellular Signaling in Cardiovascular Development and Disease Laboratory, Centro Nacional de Investigaciones Cardiovasculares Carlos III (CNIC), Calle Melchor Fernández Almagro 3, 28029 Madrid, SPAIN

[2]Ciber de Enfermedades Cardiovasculares, Instituto de Salud Carlos III, Calle Melchor Fernández Almagro 3, 28029 Madrid, SPAIN

[3]Departamento de Farmacología, Facultad de Medicina, Universidad Complutense de Madrid, Plaza Ramón y Cajal S/N, 28040 Madrid, SPAIN

[4]Unidad CSUR de Cardiopatías Familiares, Servicio de Cardiología, Hospital Universitario Virgen de la Arrixaca, Universidad de Murcia, El Palmar, 30120 Murcia, SPAIN

Corresponding author: José Luis de la Pompa


---

#### Supporting Information

**Supplementary Figures & Legends**

**Supplementary Tables and Movies legends.**

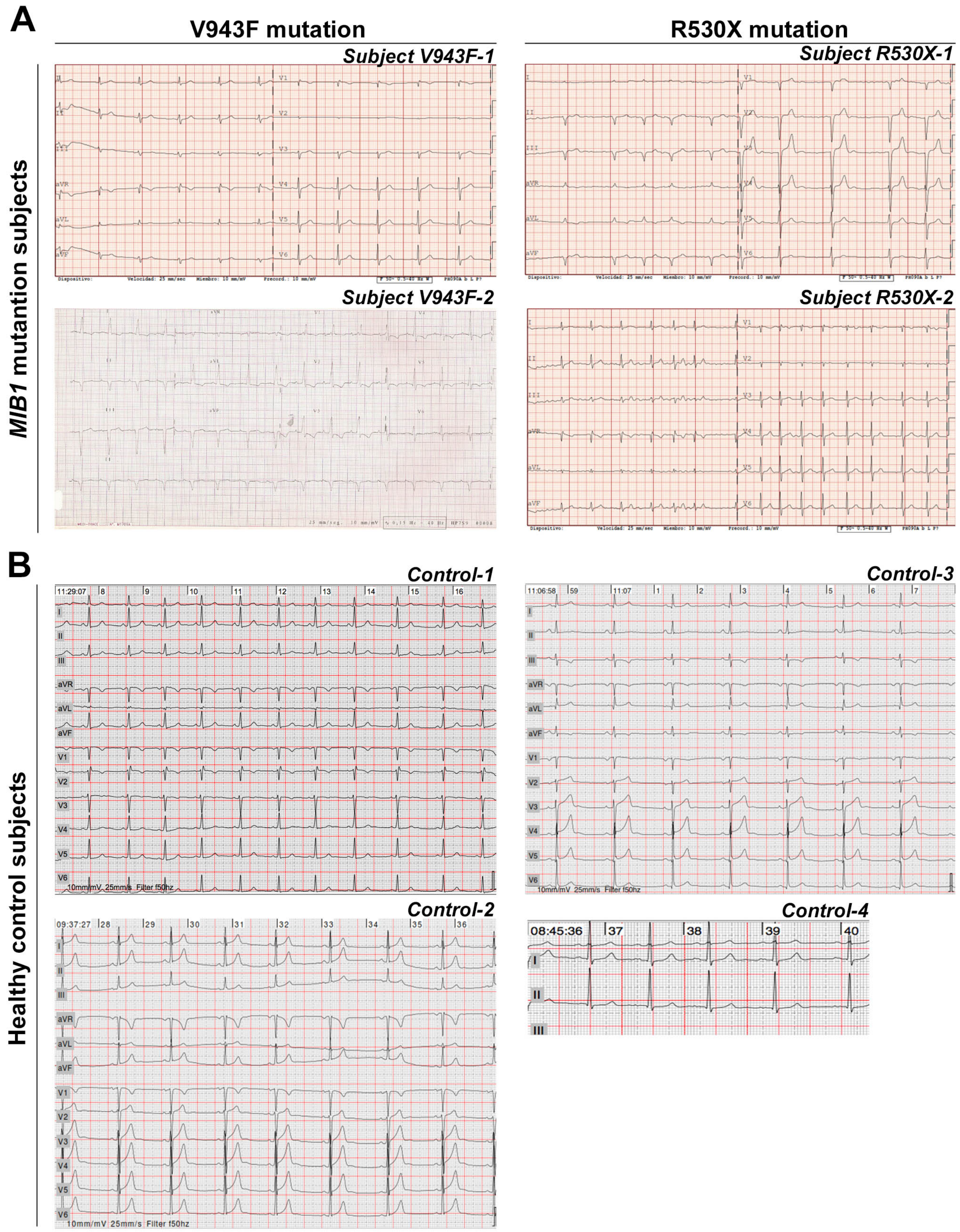

A
V943F mutation
R530X mutation
Subject V943F-1
Subject R530X-1
Subject V943F-2
Subject R530X-2
MIB1 mutantion subjects
B
Control-1
Control-3
Control-2
Control-4
Healthy control subjects


### S1 Figure

**S1 Fig. Electrocardiographic recordings from patients carrying MIB1 mutations and healthy relatives.** **(A)** Original ECG recordings from 4 individuals presenting MIB1$^{VF943F}$ (2 individuals) and MIB1$^{R530X}$ mutations (2 individuals). **(B)** Original ECG recordings from 4 healthy relatives.

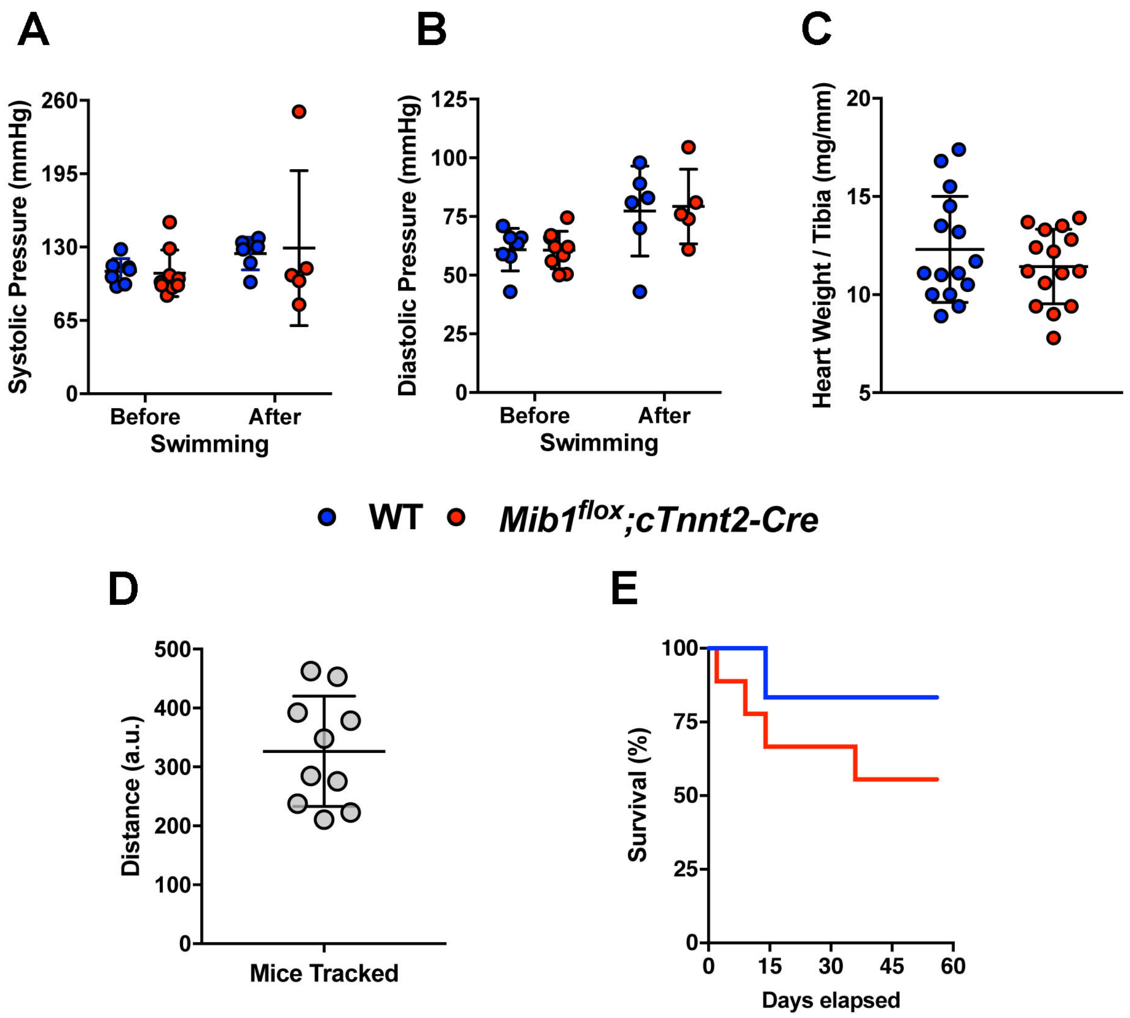

A
Systolic Pressure (mmHg)
260
195
130
65
0
Before
After
Swimming
B
Diastolic Pressure (mmHg)
125
100
75
50
25
0
Before
After
Swimming
C
Heart Weight / Tibia (mg/mm)
20
15
10
5
WT
Mib1flox;cTnnt2-Cre
D
Distance (a.u.)
500
400
300
200
100
0
Mice Tracked
E
Survival (%)
100
75
50
25
0
0
15
30
45
60
Days elapsed


### S2 Figure

**S2 Fig. Blood Pressure analysis**. Averaged systolic pressure (A) and diastolic pressure (B) of WT and $Mib1^{flox};Tnnt2^{Cre}$ mice did not show differences between groups neither before and after swimming endurance training. Statistical significance was determined by ANOVA followed by the Tukey post-hoc test for multiple comparisons. Results are expressed as mean±SD of 7-6 WT and 9-5 $Mib1^{flox};Tnnt2^{Cre}$ mice. (C) Ratio Heart Weight/Tibia. Summary data showing no differences between WT and $Mib1^{flox};Tnnt2^{Cre}$ mice. Statistical significance was determined by unpaired two-tailed Student's t-test. Results are expressed as mean±SD of 15 WT and 15 $Mib1^{flox};Tnnt2^{Cre}$ mice. (D) Homogeneity of training intensity. Analysis of the distance swum determined from consecutive time-lapse images for 1 min video recorded (online video 1 and 2, the dots and lines represent each animal and lines represent the tracking of a single animal.). The data is adjusted to a Gaussian pattern (passed the D'Agostino & Pearson normality test, alpha = 0.05). (E) Survival curve of mice during endurance training. Analysis of WT and $Mib1^{flox};Tnnt2^{Cre}$ mice survival percentage during the endurance swimming.

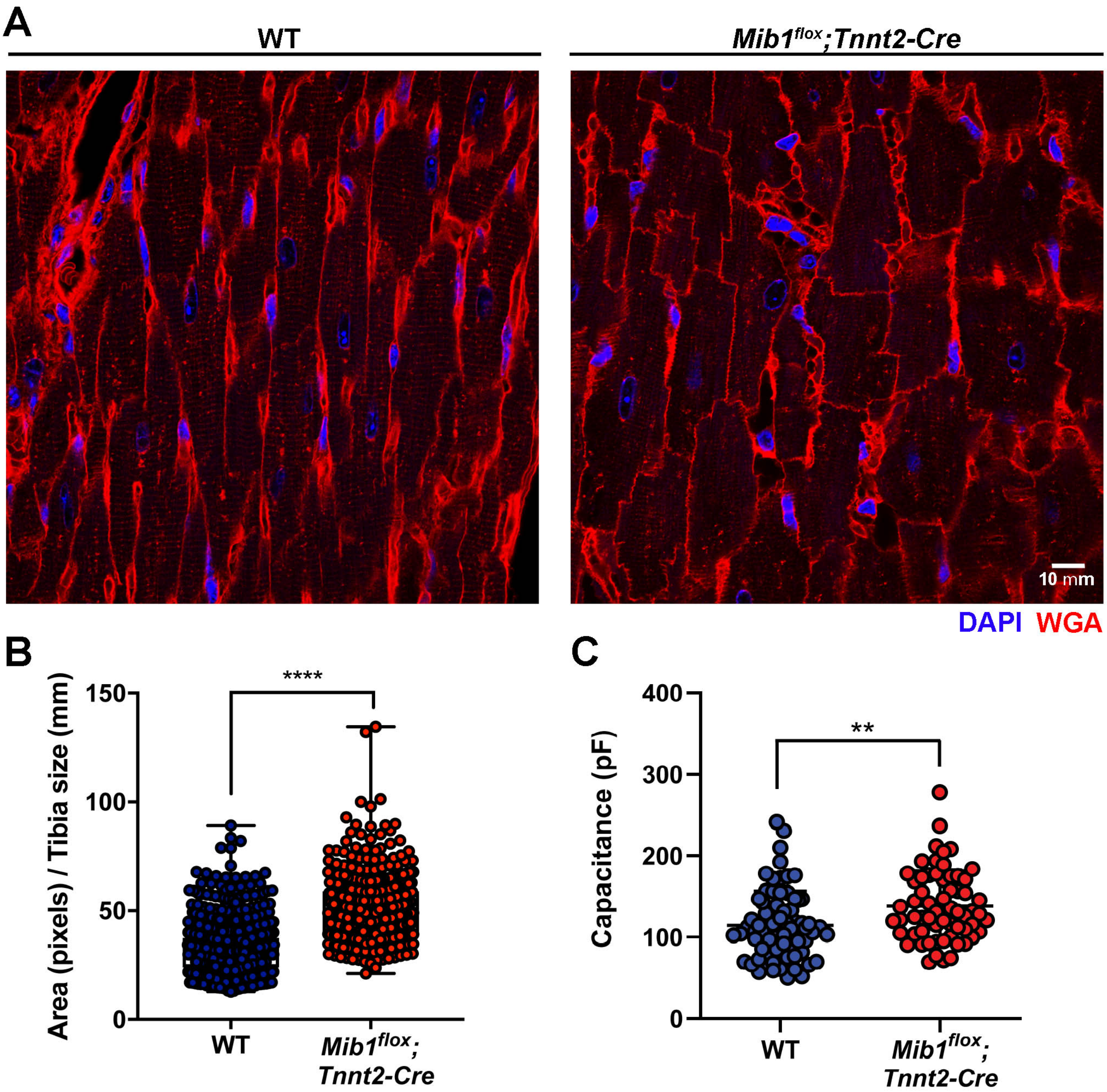
A
WT
Mib1flox;Tnnt2-Cre
10 mm
DAPI WGA
B
****
150
100
50
0
Area (pixels) / Tibia size (mm)
WT
Mib1flox; Tnnt2-Cre
C
**
400
300
200
100
0
Capacitance (pF)
WT
Mib1flox; Tnnt2-Cre

S3 Figure

**S3 Fig. Wheat germ agglutinin (WGA) staining in mouse heart sections. (A)** Representative confocal image of myocardial sections exhibiting increased cardiomyocyte size in the *$Mib1^{flox}$;$Tnnt2^{Cre}$* mice. (B) Summary data showing that cardiomyocytes area/tibial ratio is increased in *$Mib1^{flox}$;$Tnnt2^{Cre}$* mice. (C) Summary data showing that cardiomyocyte capacitance is increased in *$Mib1^{flox}$;$Tnnt2^{Cre}$* mice. Statistical significance was determined by unpaired two-tailed Student's t-test. To take into account repeated sample assessments, data were analysed with multilevel mixed-effects models. $^{**}P < 0.01$; $^{****}P < 0.0001$ vs WT. In B, results are expressed as mean±SD of 400-600 cells from 3 animals per genotype. In C results are expressed as mean±SEM of 55-60 cells from 5 animals per genotype.

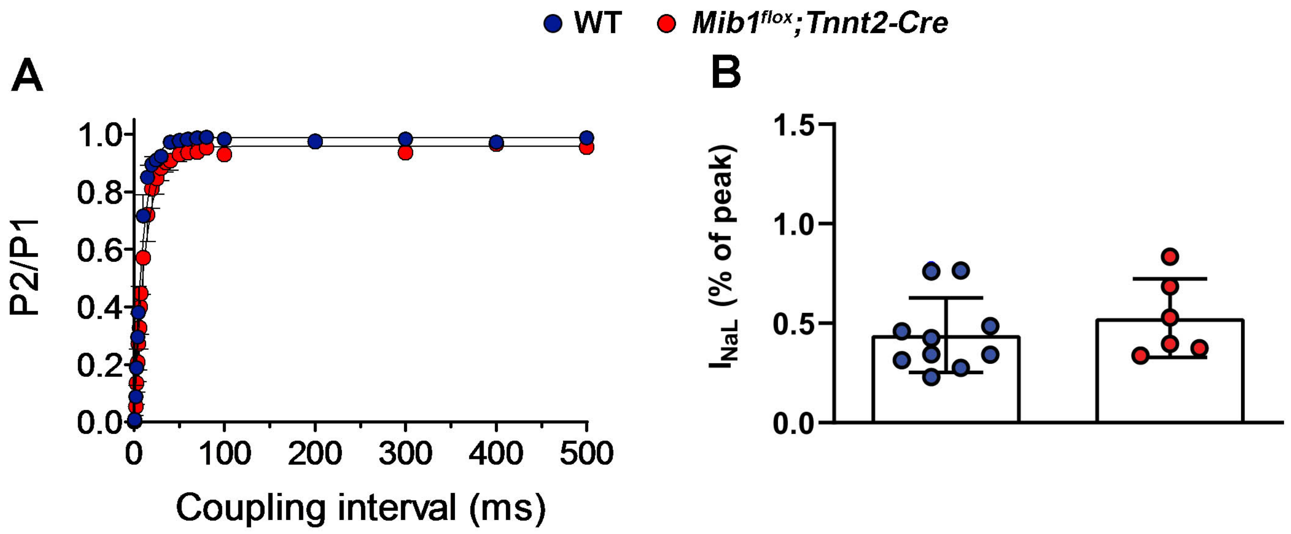
WT
Mib1flox;Tnnt2-Cre
A
P2/P1
1.0
0.8
0.6
0.4
0.2
0.0
0
100
200
300
400
500
Coupling interval (ms)
B
INaL (% of peak)
1.5
1.0
0.5
0.0

**S4 Figure**

**S4 Fig. *Mib1* inactivation did not modify $I_{Na}$ recovery from inactivation or the $I_{NaL}$.** (A) Time course of peak INa recovery from inactivation measured by using a double-pulse protocol (see supplementary methods) in WT and *Mib1$^{flox}$;Tnnt2$^{Cre}$* mice. Continuous lines represent the fit of a monoexponential function to the data. Each point represents the mean±SEM of n experiments/cells. (B) INaL (expressed as percentage of the peak current) recorded at the end of 500-ms pulses to -40 mV from a holding potential of -120 mV. Unpaired two-tailed Student's t-test was used. Statistical significance was confirmed by using non-parametric tests (two-sided Wilcoxon's test) for small-size samples (n<15). To take into account repeated sample assessments, data were analyzed with multilevel mixed-effects models. Each bar represents the mean±SEM of 10 WT and 6 *Mib1$^{flox}$;Tnnt2$^{Cre}$* cardiomyocytes dissociated from 5 animals per genotype.

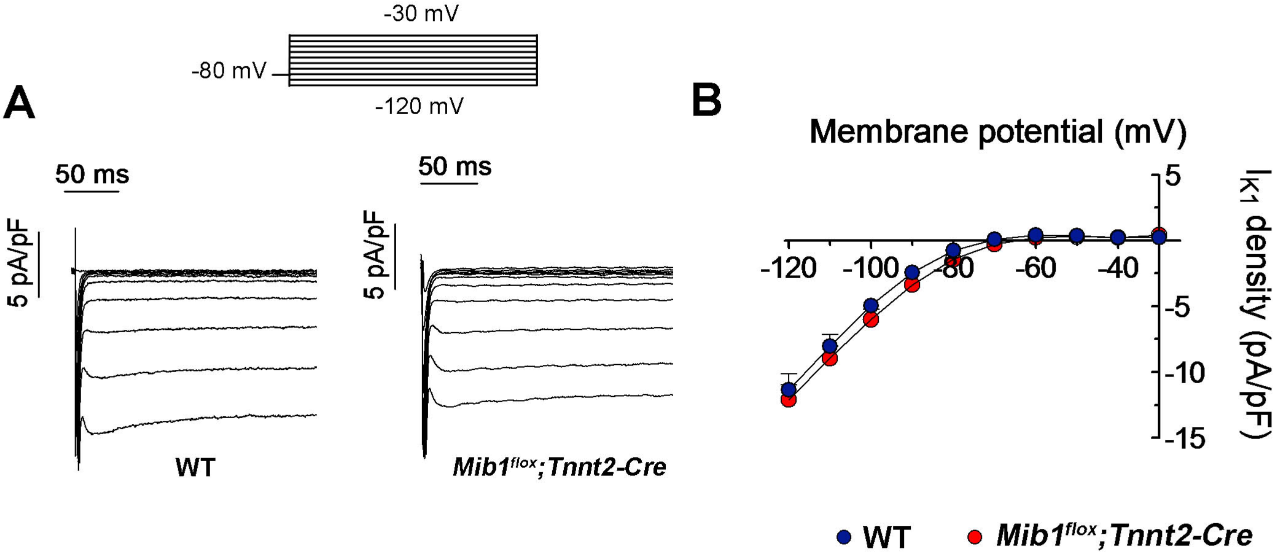
-30 mV
-80 mV
-120 mV
A
50 ms
5 pA/pF
WT
50 ms
5 pA/pF
Mib1flox;Tnnt2-Cre
B
Membrane potential (mV)
-120
-100
-80
-60
-40
5
-5
-10
-15
IK1 density (pA/pF)
WT
Mib1flox;Tnnt2-Cre

**S5 Figure**

**S5 Fig. Effects of *Mib1* inactivation on $K^+$ currents.** (A) Representative $I_{K1}$ traces recorded in two cardiomyocytes from WT or *Mib1$^{flox}$;Tnnt2$^{Cre}$* mice by applying the pulse protocol shown at the top. (B) Mean current density-voltage curves for $I_{K1}$ recorded in cardiomyocytes from both mouse groups. Unpaired two-tailed Student's t-test was used. Statistical significance was confirmed by using non-parametric tests (two-sided Wilcoxon's test) for small-size samples (n<15). To take into account repeated sample assessments, data were analysed with multilevel mixed-effects models. Results are expressed as mean±SEM of 19 WT and 14 *Mib1$^{flox}$;Tnnt2$^{Cre}$* cardiomyocytes dissociated from 5 animals per genotype.

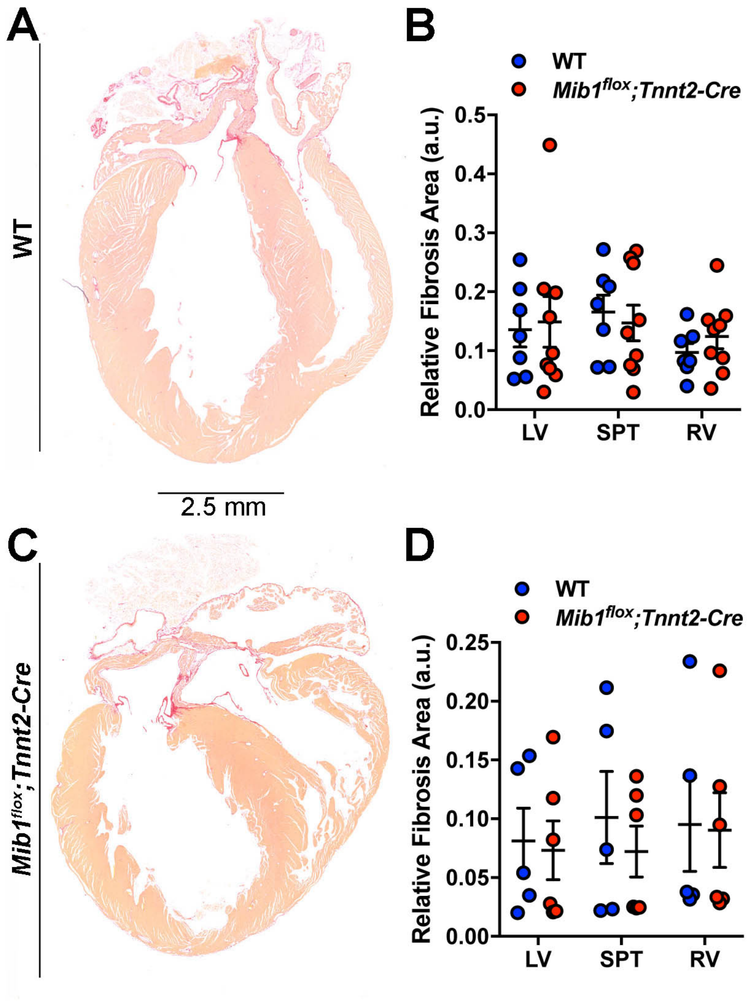
A
WT
2.5 mm
B
WT
Mib1flox;Tnnt2-Cre
Relative Fibrosis Area (a.u.)
0.5
0.4
0.3
0.2
0.1
0.0
LV
SPT
RV
C
Mib1flox;Tnnt2-Cre
D
WT
Mib1flox;Tnnt2-Cre
Relative Fibrosis Area (a.u.)
0.25
0.20
0.15
0.10
0.05
0.00
LV
SPT
RV

### S6 Figure

**S6 Fig. Picrosirius red of collagen staining in mouse heart sections.** Picrosirius red observed under bright field microscopy in WT (A) and *Mib1$^{flox}$;Tnnt2$^{Cre}$* (C) heart section did not exhibit myocardial fibrosis. (B and D) Quantification of the areas occupied by picrosirius-positive collagen in WT and *Mib1$^{flox}$;Tnnt2$^{Cre}$* mice after endurance swimming (B) and isoproterenol (D) protocol. Statistical significance was determined by ANOVA followed by the Tukey post-hoc test for multiple comparisons. In B and D, each point represents the mean of 3 section per animal. Results are expressed as mean±SD of 5-7 WT and 6-9 *Mib1$^{flox}$;Tnnt2$^{Cre}$* mice.

**Supplementary Tables and Movies legends.**

**S1 Table.** ECG parameters from patients with the *MIB1*$^{VF943F}$, *MIB*$^{1R530X}$ mutations and control.

**S2 Table.** QT variability measurements from patients with the *MIB1*$^{VF943F}$, *MIB*$^{1R530X}$ mutations and control.

**S3 Table.** Swimming endurance training protocol.

**S4 Table.** $I_{Na}$, $I_{CaL}$ and outward $K^+$ currents parameters for WT and *Mib1*$^{flox}$*;Tnnt2*$^{Cre}$ cardiomyocytes.

**S1 Video**. Video showing WT and *Mib1*$^{flox}$*;Tnnt2*$^{Cre}$ mice swimming.

**S2 Video**. Video tracing WT and *Mib1*$^{flox}$*;Tnnt2*$^{Cre}$ mice when swimming.
